# Insights into the regulation of VPS13 family bridge-like lipid transfer proteins from the structure of VPS13C

**DOI:** 10.1101/2025.11.10.687702

**Authors:** Dazhi Li, Xinbo Wang, Bodan Hu, Hongyan Hao, Stephanie Hamill, Yuting Li, Guochao Chen, Pietro De Camilli, Karin M. Reinisch

## Abstract

Bridge-like lipid transfer proteins (BLTPs) play central roles in redistributing lipids from their primary site of synthesis in the endoplasmic reticulum to other organelles. They comprise bridge-domains spanning between organelles at contact sites that allow lipids to transit the cytosol between adjacent membranes. The assembly of BLTPs into complexes with adaptor proteins enables their lipid transfer ability. To address the mechanisms underlying assembly and regulation of BLTP complexes, we used cryo-EM to resolve the structure of one such BLTP, the Parkinson’s protein VPS13C, at near-atomic resolution. The structure identifies a lipid-transfer-nonpermissive conformation, where the built-in C-terminal VAB adaptor module blocks the end of the lipid transfer bridge, interfering with lipid delivery. We also identify calmodulin, central to calcium signaling, as a VPS13 partner, suggesting calcium regulation of VPS13 function. Altogether, this structure of intact VPS13C serves as starting point to understand its regulation and, more broadly, that of other BLTPs.

## Introduction

In eukaryotic cells, the lipids comprising organellar membranes are synthesized mainly in the endoplasmic reticulum (ER) and then redistributed from there to other organelles. While vesicle trafficking is long known as a mechanism for bulk lipid transfer between organelles, the fundamental role of very large (∼200-575 kDa) rod-like proteins, so-called bridge-like lipid transport proteins (BLTPs), is only recently recognized (Hanna et al., 2023; Levine, 2022; Reinisch et al., 2025). These proteins can span between the ER and other organelles at contact sites, where the organelles are closely apposed, via “bridge” domains. These domains feature hydrophobic grooves that allow lipids to travel through the cytosol between organellar membranes. Lipid transfer via BLTP underlies multiple functions, including membrane expansion, as required for the biogenesis of new organelles such as the autophagosome, or the maintenance of other organelles, such as mitochondria and peroxisomes which are disconnected from vesicle trafficking pathways and so rely exclusively on protein-mediated transport for their membrane lipid supply.

Several recent studies have shown that BLTPs do not work alone but rather in complexes with partner proteins (Dziurdzik and Conibear, 2021; Hanna et al., 2023; Reinisch et al., 2025) that can include adaptors to anchor the tips of the BLTP bridge domain to donor and acceptor organelles, as well as integral membrane proteins that facilitate the transfer or redistribution of lipids at the BLTP-membrane junction. Identification of partner proteins and of the mechanisms that control their assembly with VPS13, including potential conformational changes of VPS13, are subject of ongoing investigations and are prerequisite for understanding the molecular mechanisms underlying BLTP-mediated lipid transfer and its regulation.

As founding members of the BLTP super family, VPS13 proteins are among the best studied, including by our groups(Hanna et al., 2023; Reinisch et al., 2025). They are conserved across eukaryotes, with a single protein in yeasts and four versions (A-D) in humans. The VPS13s feature built-in adaptor modules at the C-terminal end of the bridge domain, which mediate their association with the lipid acceptor organelle. These include amphipathic helices in the ATG2_C region (so-called because such helices are also present in ATG2 proteins, another BLTP family) and the PH-and VAB domains as well as a WWE-domain in VPS13A and -C (Fig1A). In human VPS13s (except VPS13B), a peptide (called “FFAT”) motif near the N-terminal end of the bridge-domain mediates their association with the lipid donor organelle, the ER, via an interaction with the ER-resident VAP (Kumar et al., 2018). The identity of other proteins that might couple the N-terminal end of VPS13 to the donor organelle had heretofore been elusive.

In humans, VPS13 dysfunction is associated with severe neurological diseases (Hanna et al., 2023), including chorea acanthocytosis for VPS13A (Rampoldi et al., 2001; Ueno et al., 2001) and an early onset version of Parkinson’s for VPS13C (Lesage et al., 2016), making them of significant biomedical interest. Whereas the single yeast VPS13 multi-tasks, each of the human VPS13’s has distinct localizations and functions in the cell (Hanna et al., 2023), explaining their different disease manifestations. VPS13s play roles in mitochondrial maintenance; in the formation of autophagosome-, prospore (yeast)-, and acrosomal membranes; in peroxisome biogenesis; and in Golgi homeostasis (Reinisch et al., 2025). VPS13C, in particular, plays a key role in lysosome homeostasis as it is localized at contacts between the ER and late endosomes and lysosomes (hence referred to as lysosomes). This localization is highly regulated as VPS13C is massively recruited to these contacts in response to lysosomal damage (Wang et al., 2025), presumably to deliver lipids for repair of the lysosome membrane.

Although first insights into the structure of VPS13 proteins were obtained from x-ray and cryo-electron microscopy (cryo-EM) studies of VPS13 fragments (Kumar et al., 2018; Li et al., 2020) and complemented by Alphafold predictions (Levine, 2022), our understanding of their mechanism of action and regulation has been hindered by the lack of experimental structural information for any intact protein, especially as it pertains to the disposition of the adaptor domains with respect to the bridge domain. Here, to obtain such insights, we used cryo-EM to visualize an intact VPS13, the Parkinson’s protein VPS13C (425 kDa), at near atomic resolution (in 3 maps, 3.8-4.2 Å). The maps featured density close to the N-terminal tip of the bridge domain that could not be accounted for by VPS13C alone, and which we subsequently identified as belonging to calmodulin (CaM), which co-purifies with VPS13s in a high affinity complex. This finding raises the possibility that CaM may participate in controlling VPS13 recruitment and/or attachment to the lipid donor membrane and, moreover, highlights growing evidence for calcium signaling in regulating VPS13 activity. At the other, C-terminal end of VPS13C, the reconstruction shows the VAB domain arching over the bridge-domain to interfere with its binding to the acceptor membrane and hence with lipid transfer. Thus, purified VPS13C, likely representing the cytosolic form of VPS13C not engaged at membrane contact sites, is in a lipid-transfer incompetent state. The protein must undergo conformational changes as it engages with the acceptor membrane to allow the bridge domain to access this membrane. As the VAB domain is a common feature of all VPS13s, they likely all feature lipid transfer inactive and active conformations, and the transition between conformations offers means for regulating their activity.

## Results

### Cryo-EM Reconstruction

We undertook biochemical and single particle cryo-EM studies to better understand how the function of VPS13, and particularly of VPS13C, may be regulated. To this aim, we overexpressed a FLAG-tagged version of VPS13C (VPS13C-3XFLAG) in Expi293 cells and isolated it using anti-FLAG resin affinity purification, followed by size exclusion chromatography (Suppl. Table S1, Suppl. Fig. S1).

We collected 9054 electron micrographs, which we processed in CryoSPARC (Punjani et al., 2017) to generate 3 maps (Fig. 1B-C, Sup. Fig. S2). An initial map (#1) shows intact VPS13C at a nominal resolution of 4.2 Å. After template matching and rounds of refinement, we obtained a locally-refined map (#2) at a nominal resolution of 3.8 Å, showing the C-terminal portions of VPS13C. A separate reconstruction (map #3) allowed us to visualize N-terminal portions of VPS13C at a nominal resolution of 4.1 Å. We used an Alphafold (Jumper et al., 2021) generated version of VPS13C to guide model building into the maps. Alphafold predicts that the bridge domain comprises a series of 13 repeating beta groove (RBG) motifs, each consisting of 5 antiparallel beta strands, arrayed end-to-end into a taco shell shaped structure, with “caps” at the very N- and C-terminal tips of the bridge domain. We dissected the bridge-domain into RBG and “cap” segments, and placed these into the maps, then adjusted the model manually. Similarly, we placed Alphafold models for the VAB-, PH- and WWE domains as rigid bodies, followed by manual rebuilding when appropriate. Well resolved density in map #3 allowed modeling for the N-terminal cap and RBGs 1-5, and map #2 allowed modeling of RBGs 8-13, the C-terminal “cap” and the VAB, WWE, and PH domains. RBGs 6-7 were modeled based on Map#1. There was no density corresponding to the amphipathic helices in the ATG2_C motif in any of the maps, suggesting they are unstructured, mobile, or both, in the absence of membrane. Map #3 showed density that could not be accounted for by the AlphaFold model for VPS13C, spanning over the taco shell between RBGs 1 and 2, and this density was subsequently modeled as CaM (Fig. 1C, see below).

**Figure 1.**
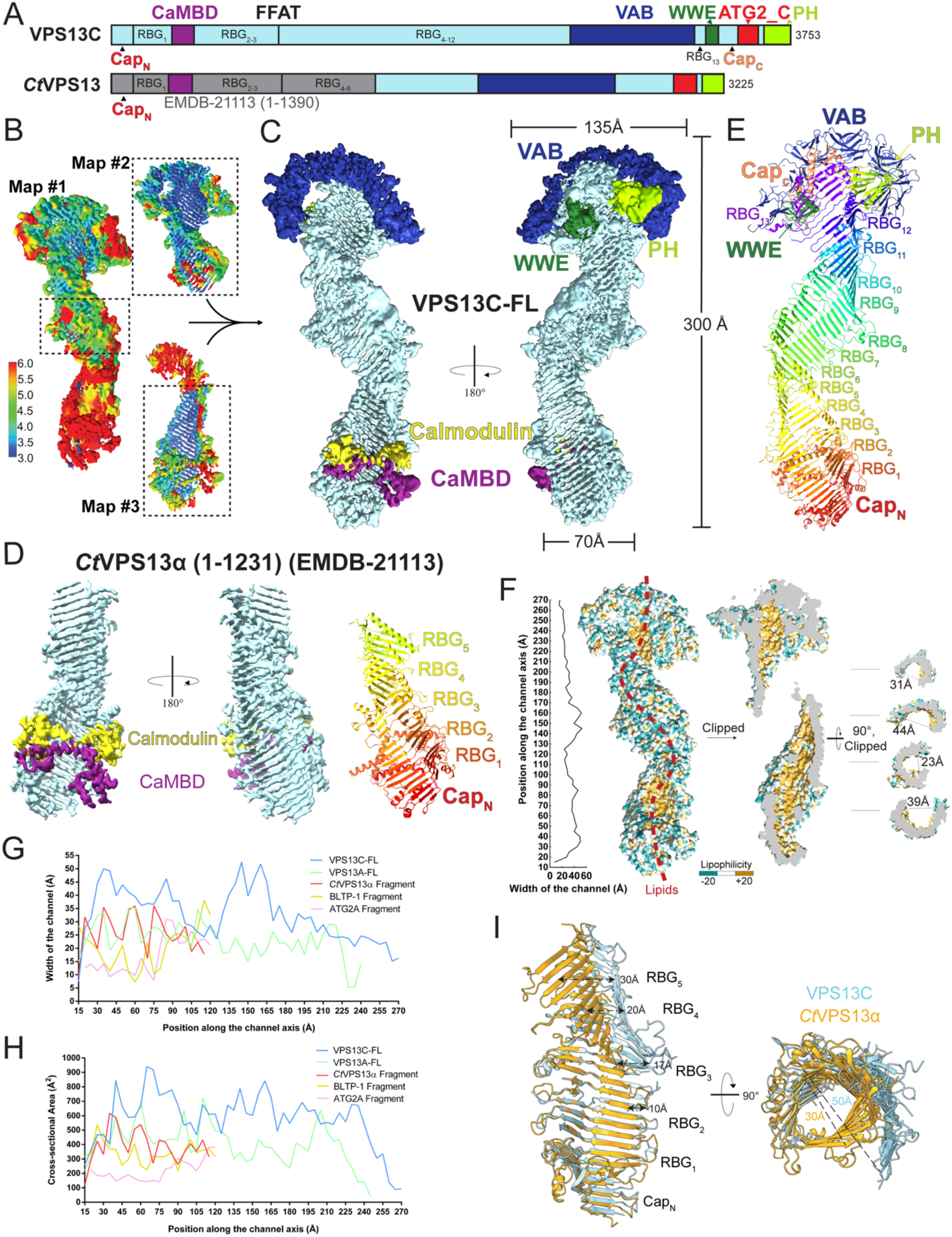
Structural analysis of VPS13C and *Ct*VPS13α in complex with CaM. (**A**) Domain architecture of human VPS13C and *Ct*VPS13α. (**B**) For VPS13C, two locally-refined maps (#2 and #3) were aligned and combined with the central portion of the full-length map (#1) to generate a composite map. Maps are colored according to local resolution (also see Suppl. Fig. S3 and Suppl. Table S2.). (**C**) A composite map for the VPS13C/CaM complex is shown, with the bridge domain colored in light blue. The C-terminus of VPS13C, consisting of the VAB domain (dark blue), the PH domain (light green), and the WWE domain (dark green) contacts the lysosome membrane at contact sites. VPS13C’s N-terminus interacts with CaM (yellow) through a ‘handle-like’ region, referred to as CaM binding domain (CaMBD, in purple). (**D**) Cryo-EM map of the N-terminal fragment of *Ct*VPS13α (EMDB-21113) showing the CaMBD (purple) and copurified CaM (yellow). The right panel shows the model for *Ct*VPS13α (1-1231). The N-terminal “cap” and five individual RBG domains are shown in different colors. (**E**) Model of VPS13C alone, with different colors for the terminal “caps”, thirteen RBG domains, and adaptor domains. Dashed lines indicate flexible loops unresolved in the EM maps. (**F**) VPS13C harbors a hydrophobic channel with varying width. Representative cross-sections are on the right. The red dotted line indicates the presumed lipid transfer path. (**G, H**) Comparison of the channel width and the cross-sectional area along the channel axis in different experimental structures of BLTPs, calculated as for (F). VPS13 family proteins, especially VPS13C, exhibit a wider channel. (**I**) Overlay of *Ct*VPS13α with the N-terminal fragment of VPS13C containing the N-terminal cap and the first five RBG motifs. For clarity, some connecting segments between the RBG ß-strands are omitted.

Guided by Alphafold (Jumper et al., 2021), we also modeled N-terminal portions of VPS13 from the fungus *Chaetomium thermophilum* (residues 2-1230), based on a previously reported single particle reconstruction at 3.8 Å nominal resolution (EMD #21113, (Li et al., 2020)). Similarly to our VPS13C sample, the fungal VPS13 fragment used in the reconstruction, which we refer to as *Ct*Vps13α, was purified from Expi293 cells, and we found density corresponding to CaM in this reconstruction as well (Fig. 1D). The CaM binding site is equivalent in VPS13C and the fungal Vps13.

Suppl. Figure S3 and Suppl. Table S2 show representative portions of the maps and statistics for data collection, processing, and modeling.

### Architectural overview of the VPS13C-CaM complex

At low resolution, VPS13C resembles a bubble wand, with the bridge-domain as a ∼ 300 Å long handle and the C-shaped VAB domain as the loop. As predicted by AlphaFold, the bridge domain comprises N- and C-terminal “caps” at either end of the handle, with RBG motifs arrayed in between (Fig. 1 C, E). The VAB is an insertion between RBGs 12 and 13, and is positioned over the C-terminal end of the bridge domain by interactions with linkers in the C-terminal cap. The WWE and PH domains are positioned along the convex back face of the taco shell. Calmodulin is bound at the N-terminal end of the bridge domain (Fig. 1C).

### Bridge domain

Residues on the convex solvent exposed exterior of the bridge domain’s taco shell are hydrophilic, whereas residues lining the interior comprising the lipid transport path are hydrophobic (Fig. 1F). Lipid fatty acyl moieties are expected to bind within the hydrophobic channel, with their headgroups exposed to solvent. The channel appears filled at lower contour levels (not shown), but we could not resolve individual lipids. This is due in part to the limited resolution of the reconstructions and also because each reconstruction represents the average of hundreds of thousands of particles, where lipids do not necessarily have well defined binding sites. We estimate that the channel can accommodate ∼120 lipids. The channel’s width varies along the length of the bridge domain: it measures ∼20 Å across at its narrowest and ∼50 Å across at its broadest points.

In addition to VPS13C and *Ct*Vps13α modeled by us here (Fig. 1D-E), several other experiment-based structures have recently become available for other BLTPs: VPS13A (Hu et al, in preparation), ATG2 (Wang et al., 2024), and an N-terminal fragment of BLTP-1 (Kang et al., 2025). The width of the lipid transfer channel is variable along the length of the bridge domain in all of these proteins. Whether this variability is physiologically significant is unclear. Notably, however, the lipid transfer channel is wider in all of the VPS13 proteins compared to either ATG2 or BLTP1 (Fig. 1G-H), and it is significantly wider in VPS1C even as compared to VPS13A (Hu et al, in preparation) or fungal VPS13 (Fig. 1G-I). We posit that the width of the lipid transfer channel could impact lipid flux, so that the VPS13s, and VPS13C in particular, may be able to transfer lipids at faster rates than other narrower BLTPs. Lending plausibility, a recent manuscript reports that mutations that constrict ATG2’s width affect lipid transfer rates in cells (bioRxiv 2025.05.24.655882).

### CaM is a VPS13 interaction partner at the N-terminal end of the VPS13 bridge-domain

Mass spectrometry analysis of proteins that co-immunoprecipitated with overexpressed VPS13C from Expi293F cell extracts identified CaM as a major hit (Suppl. Table S3). We confirmed that CaM was present in the human VPS13C sample we purified for structural studies by western blotting (Fig. 2A). CaM is also reported as an interactor of VPS13A, C and D, but not of VPS13B (the most divergent member of the VPS13 family), in proteomics databases (BioGrid) and was detected by western blotting of VPS13A purified from Expi293 cells (Hu et al, in preparation). Moreover, an N-terminal fragment of VPS13D was identified in a screen for CaM interactors in *C.elegans* (Shen et al., 2008). Consistent with our findings, AlphaFold (Abramson et al., 2024) predicts with high confidence a complex between CaM and human VPS13A, C, and D, but not VPS13B, and in these predictions CaM is positioned where the previously unexplained density in our maps of the VPS13C N-terminus was localized (map #3) (Suppl. Fig. S4). Accordingly, we modeled CaM into this density (Fig. 2B). The interfaces between CaM and VPS13s are large (total buried surfaces between CaM and VPS13C and VPS13A, respectively, are 4380 A^2^ and 5713 A^2^), indicative of a high affinity interaction (Yan et al., 2008). In the VPS13C reconstruction, CaM binds to four helices (H_334-355_ comprising residues 334-355, H_370-391_ comprising residues 370-391, H_395-418_ comprising residues 395-418, and H_438-451_ comprising residues 438-451) that span across the bridge domain to connect the first and second RBG (Fig. 2B). CaM binds equivalent helical elements in another experimental structure, the bridge domain of VPS13A (Hu et al, in preparation). VPS13B, which does not appear to bind calmodulin based on proteomics databases (BioGRid), lacks the series of helices that form the primary calmodulin binding site (Suppl. Fig S4).

**Figure 2.**
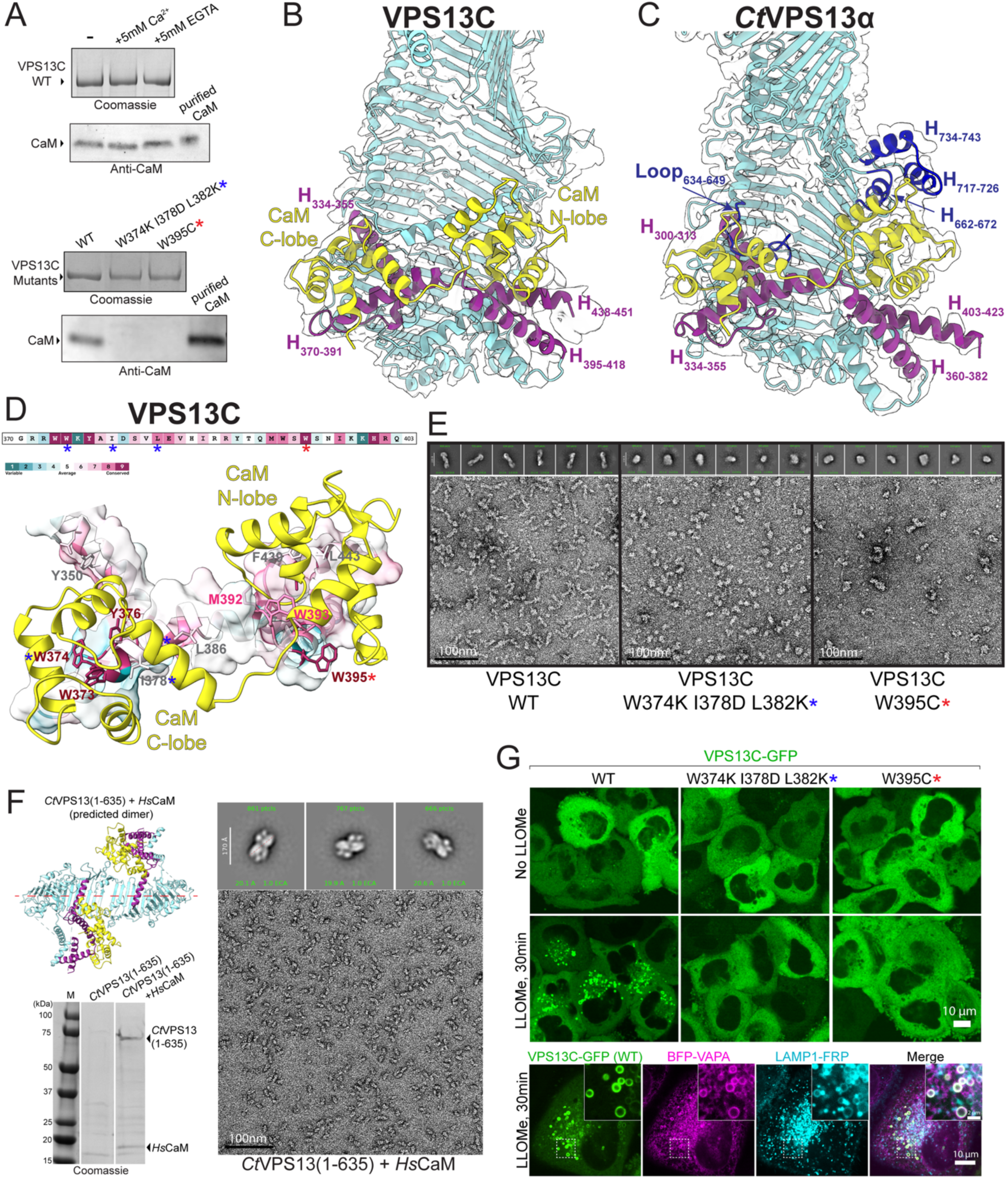
CaM is a constitutive VPS13 binding partner at the N-terminus. (**A**) Co-purified CaM in FLAG-IP, detected by western blot. CaM association with VPS13C is unaffected by purification in the presence of 5mM Ca^2+^ or 5 mM EGTA. However, because substrate binding can raise CaM’s calcium affinity by over 1000-fold, enabling binding at very low calcium concentrations in the presence of EGTA (Jurado et al., 1999), we cannot exclude that the VPS13C-CaM interaction is sensitive to calcium levels. The bottom panel shows that the interaction is completely abolished when mutations are introduced to disrupt the interaction with either the N-terminal or C-terminal lobe of CaM. Blue and red stars indicate the W374K/I378D/L382K mutant and the W395C mutant, respectively. Because of the difference in sizes for VPS13C and CaM, the same samples were loaded onto two different gels for analysis: 3-8% Tris-acetate to better resolve VPS13C and 4-20% Tris-glycine to better resolve CaM. (**B**) At VPS13C’s N-terminus, CaM’s N-terminal lobe interacts with H_395-418_ and H_438-451_, while the C-terminal lobe interacts with H_334-355_ and H_370-391_. All four helices are positioned between RBG_1_ and RBG_2_. (**C**) At *Ct*VPS13’s N-terminus, the N-terminal lobe of CaM interacts with H_360-382_ and H_403-423_, while the C-terminal lobe interacts with H_300-313_ and H_334-355_. Additionally, Loop_634-649_, H_662-672_, H_717-726_, and H_734-743_ from between RBG_2_ and RBG_3_ contact CaM near its EF-hand loops. Equivalent regions are absent from the VPS13C structure. (**D**) Conservation scores for VPS13C calculated through the ConSurf web server (Ashkenazy et al., 2016). The bottom panel shows an enlarged view of the conserved hydrophobic residues of VPS13C interacting with the cavities in CaM’s C-and N-terminal lobes. (**E**) The representative negative-stain EM image and 2D averages showing that both the W374K/I378D/L382K mutant and the W395C mutant do not adopt WT VPS13’s rod-like shape, although they do not aggregate. (**F**) When expressed in *E. coli*, *Ct*VPS13 (residues 1-635) is insoluble unless co-expressed with CaM. The representative EM image and 2D averages show that the resulting complex forms a homogenous, end-to-end dimerized tube, consistent with AlphaFold3 prediction (colored as in previous panels). (G) After LLOMe-induced lysosome damage, wildtype VPS13C (WT) forms punctae at ER-lysosome contact sites (bottom panels). In contrast, both the W374K/I378D/L382K and the W395C mutants remain dispersed in the cytosol.

CaM comprises an N- and a C-terminal lobe, each of which can interact with substrate proteins. Each lobe has two EF-hands, which are helix-turn-helix motifs with calcium binding sites in the loop between helices (Andrews et al., 2020). In the structures of the VPS13-CaM complex, both lobes engage with VPS13. The N-terminal lobe interacts primarily with the VPS13C H_395-418_, and the C-terminal lobe interacts primarily with the VPS13C H_370-391_. VPS13 residues at the interface are conserved and hydrophobic, interacting with hydrophobic surface cavities on CaM (Fig. 2D).

We confirmed by western blotting that CaM also co-purifies with *Ct*Vps13α recombinantly produced in Expi293 cells (Suppl. Fig. S4), and we found density corresponding to CaM in the map for *Ct*VPS13 (Fig. 1D, 2C). In a previous study (Li et al., 2020), we had described this density as part of a “handle” that arches over the “basket”-shaped bridge domain and had incorrectly attributed it to Vps13. The interface area is even larger than that of the human VPS13s and CaM (6052 A^2^ total buried surface area). As in the VPS13C and VPS13A cryo-EM structures, CaM interacts with helices that span between RBGs 1 and 2 (*Ct*H_300-313_, *Ct*H_334-355_, *Ct*H_360-382,_ *Ct*H_403-423_). Additionally, the calcium binding loop of the N-terminal lobe’s second EF-hand interfaces with a 3-helix bundle formed from loops of Vps13’s third RBG repeat (*Ct*H_662-672_, *Ct*H_717-726_, *Ct*H_734-743_). A similar interaction is present in the VPS13A structure (Hu et al, in preparation), but is not present in the VPS13C maps. Finally, the C-terminal lobe of CaM interacts with a loop in *Ct*Vps13’s second RBG repeat (residues 634-649). Note that endogenous yeast Vps13 was found to co-purify with Cdc31, a member of the CaM superfamily (De et al., 2017; Kilmartin, 2003). Based on AlphaFold prediction, Cdc31 binds where CaM is bound in the cryo-EM reconstruction of recombinant *Ct*Vps13 (Supp. Fig. S4). We note that other members of the CaM superfamily, namely S100 proteins (Suppl. Table S3), also co-immunoprecipitated with VPS13C, although not with the same enrichment as CaM. Thus, the interaction between VPS13s (except VPS13B) and proteins in the CaM superfamily is conserved across eukaryotes.

To further confirm the importance of the helical segment between VPS13 RBG repeat 1 and RBG repeat 2 for CaM binding, we introduced mutations in this region of VPS13C, expecting to abrogate its interaction with CaM. We mutated residues along H_370-391_ or H_395-418_ from hydrophobic to hydrophilic to disrupt binding to CaM’s C-terminal or N-terminal lobe (Fig. 2A, D), respectively. These constructs were expressed at levels similar to the wild-type protein in our Expi293 overexpression system and did not form aggregates, such as would result from severe misfolding, as assessed by size exclusion chromatography (ie, they were not in the void volume) or negative stain EM (Fig. 2A, E, Suppl. Fig. S5). Consistent with a role for the helical segment linking RBG 1 and RBG 2 in binding CaM, the mutants did not co-purify with CaM (Fig. 2A). Interestingly, though, the mutant proteins had different profiles by size exclusion chromatography indicative of different hydrodynamic radii, and they did not form rods like wild-type protein as determined by negative stain microscopy (Fig. 2E, Suppl. Fig. S5).

These data suggest either that the mutations we introduced disrupted protein folding or else that CaM is a constitutive binding partner for VPS13 and essential for its folding. As CaM is present and highly expressed in the Expi293 cells we used for protein production, we turned to a bacterial expression system, where CaM is not expressed, to assess whether wild-type VPS13 constructs require CaM for folding. We worked with a 70kDa fragment of *Ct*Vps13 (residues 1-635), small enough for bacterial expression, that includes the tandem helices between RBG 1 and RBG 2 that comprise the primary CaM binding site. While *Ct*Vps13_1-635_ by itself was poorly expressed and not soluble, we were able to co-express it with CaM, obtaining good yields of a *Ct*Vps13_1-635_-CaM complex (Fig. 2F). *Ct*Vps13_1-635_ and CaM assemble into a heterotetramer, with the two copies of CtVps13_1-635_ arranged tail-to-tail into a rod. This tail-to-tail arrangement was also observed in previous experimental structures of *Ct*Vps13 fragments (Kumar et al., 2018; Li et al., 2020), and is predicted by AlphaFold for *Ct*Vps13_1-635_ (Fig. 2F). Thus, this experiment supports that CaM, or perhaps a member of the calmodulin superfamily for fungal Vps13, is constitutively associated with VPS13s and is required for VPS13s (except presumably VPS13B) to fold into a rod-like shape.

The location of CaM’s binding site near the N-terminal end of VPS13s, where they interact with the donor organelle, i.e. the ER (Kumar et al., 2018), raised the possibility of a role for CaM in VPS13 localization. Under control conditions the bulk of WT VPS13C is soluble in the cytosol with only a small pool of VPS13C localizing at ER-lysosome contacts. However, damage of the lysosome membrane, for example in response to addition of the lysosomotropic compound LLOMe (Wang et al., 2025), induces a massive redistribution of WT VPS13C to these organelles with the formation of ER-lysosome contacts (Fig. 2G). In contrast, consistent with a role for CaM in localizing VPS13C, we found that CaM-binding deficient mutants remain in the cytosol and fail to be recruited to either the lysosome or the ER (Fig. 2G).

### Interactions with the C-terminal “cap” stabilize the VAB in a lipid transfer inactive conformation

In the cryo-EM reconstruction of VPS13C we see that the six beta-sandwich repeats that comprise the VAB domain arch over the end of the bridge domain, positioned via extensive interactions with the C-terminal “cap” (3507 Å^2^ occluded surface)(Fig. 1E, 3A-C). This “cap” (in orange in Fig. 1E) comprises two beta strands and an alpha helix (H_3394-3418_) that lies across the bridge-domain, partially closing off the lipid transfer channel. The segment connecting RBG 13 and the cap (residues 3289-3322) snakes along the concave surface of the C-shaped VAB, contacting beta-sandwich repeats #2-#5, and the connecting loop between the cap’s two beta strands contacts sandwich repeat #4 (Fig. 3C). Additionally, in VPS13C, the end of the helix C-terminal to the PH domain (H_3731-3745_) butts against beta-sandwich repeat #1 (Fig. 3B-C). For lipid transfer to occur, both the VAB and H_3394-3418_ need move away, allowing the bridge domain access to membrane and unblocking the transfer channel.

**Figure 3.**
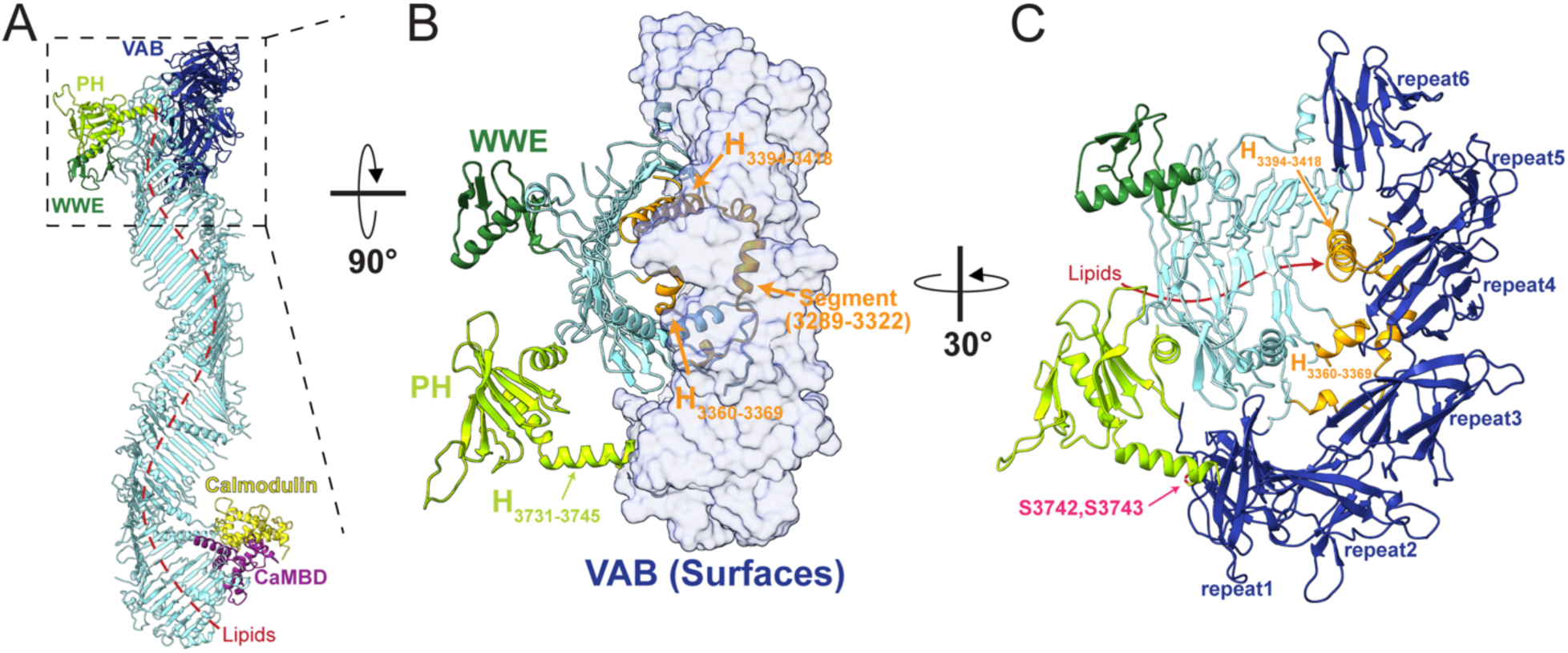
VPS13C adopts a lipid transfer-incompetent conformation at the C-terminus. (**A**) Side view of the VPS13C structure showing the VAB domain (dark blue) covering one end of the lipid transfer channel, preventing membrane docking and lipid transfer. The red dotted line indicates the presumed lipid transfer path. (**B**) VPS13C C-terminal end as viewed from the lysosome membrane. The VAB domain (pale blue surface) makes extensive contacts with the C-terminal cap (orange) through H_3418-3394_, H_3360-3369_, and an extended segment (3289-3322). At the end of the PH domain, H_3731-3745_ protrudes toward the VAB domain. (C) The C-terminal cap clamps onto beta-sandwich repeat #3 and #4 in the VAB. The presumed lipid transfer route (red arrow) is blocked by H_3418-3394_, H_3360-3369_, and the VAB domain. Additionally, H_3731-3745_ from the PH domain contacts beta-sandwich repeat #1. Phosphorylation at C-terminal residues in H_3731-3745_ (S3742, S3743), highlighted in pink, may contribute to conformational changes in the VAB domain that allow the bridge domain to access membrane.

In the cryo-EM study of another VPS13, VPS13A, in complex with a receptor at the acceptor membrane XK (Hu et al, in preparation), the VAB has moved to the side of the bridge domain, and segments of the cap that interact with the VAB in the VPS13C structure are not visible in the maps, presumably because they are mobile. Nor is there an alpha helix corresponding to VPS13C’s H_3394-3418_ blocking the end of the lipid transfer channel. Our VPS13C model here represents a lipid-transfer nonpermissive conformation, but very likely VPS13C can undergo conformational changes to a lipid transfer active conformation represented in the VPS13A structure reported by Hu *et al* (in preparation).

### VPS13C recruitment to damaged lysosomes

Previous *in cellulo* studies showed that (i) VPS13C is localized at contacts between the ER and lysosomes (Kumar et al., 2018) (Fig. 2G), (ii) that this recruitment depends on VPS13’s VAB domain and the lysosomal small GTPase Rab7 (Gillingham et al., 2019; Wang et al., 2025) and (iii) that recruitment is strikingly enhanced by lysosome damage (Wang et al., 2025). It was not known, however, whether VPS13C and Rab7 interact directly, or how.

We undertook protein-protein interaction experiments to obtain a better understanding of how VPS13C and Rab7 might bind each other (Fig. 4). In these experiments, we co-transfected mammalian cells (EXpi293 cells) with plasmids to express strep-tagged Rab7 (to ensure the GTP-bound “active” form that would be present at the lysosome, we used a constitutively active mutant, Rab7(Q67L)) and FLAG-tagged VPS13C or mutant forms of VPS13C (Fig. 4A), passed cell lysates over Strep-Tactin resin to retain Rab7(Q67L) and associated proteins, washed the resin to remove non-specifically bound proteins, and finally eluted Rab7(Q67L) and associated proteins using biotin. The eluted proteins were analyzed by SDS-PAGE and visualized by Coomassie stain and/or western blotting. Expression of Flag-tagged VPS13C constructs in the original lysates was assessed by anti-FLAG immunoprecipitation (Fig. 4B, left two panels).

**Figure 4.**
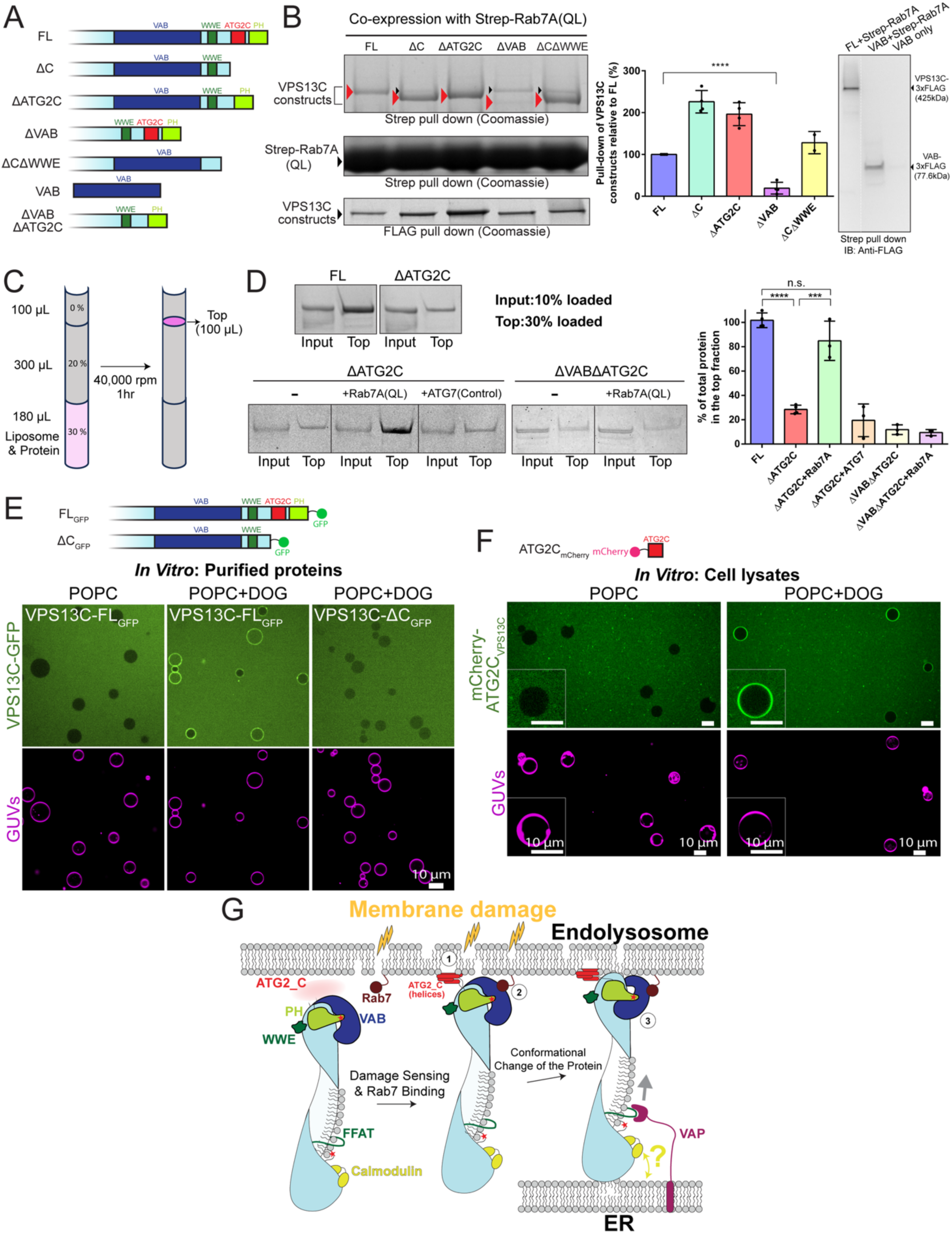
Regulatory mechanisms of VPS13C membrane recruitment. (**A**) Domain architecture of VPS13C constructs. (**B**) Strep-tagged Rab7A (Q67L) pulls down full length (FL) FLAG-tagged VPS13C when co-expressed in cells, and this interaction requires the VAB domain but not the PH domain, WWE domain, or ATG2_C region. Among all deletion constructs, only the VAB domain deletion construct drastically reduces co-purification. Red arrowheads indicate VPS13C constructs, and black arrowheads represent endogenous VPS13C. The Strep pull-down samples were analyzed by 3-8% Tris-Acetate gel, and the Flag pull-down samples were analyzed by 4-20% Tris-glycine gel. The bottom strip of the left panel shows anti-FLAG IPs of the starting lysate to indicate expression levels of the constructs. Quantification of VPS13C constructs retained by Rab7 is shown in the middle panel. The band intensity of each VPS13C construct retained by Rab7 was normalized by the corresponding Rab7 band intensity, and then normalized to the full-length (FL) construct. N=4, except ΔC+WWE, where N=2. Data were shown as mean ± s.d. and compared using a two-sided Student’s t-test; \*\*\*\**P*<0.0001. The right panel shows retention by Rab7A (Q67L) of both FL VPS13C and its VAB domain, detected by western blot. (**C**) Schematic of the liposome floatation assay. After mixed with small unilamellar vesicles (SUVs), VPS13C co-floatation correlates with its membrane binding ability. For analysis, 10% of input and 30% of the top 100 μL were loaded. (**D**) Close to 100% of VPS13C-FL and ∼30% of VPS13C-ΔATG2C co-floats with SUVs, indicating the ATG2_C region’s role in membrane binding. Recruitment of VPS13C-ΔATG2C is restored (>80% co-floatation) by tethering His-tagged Rab7A, but not the negative control His-tagged ATG7, to SUVs. Flotation of the VPS13C-ΔVABΔATG2C construct was not rescued by Rab7A tethering, indicating that recruitment of VPS13C to SUVs is promoted by the interaction between the VAB domain and Rab7. The graph (right) quantifies the fraction of VPS13C constructs that co-floated with liposomes; N=4 for FL and ΔATG2C alone; N=3 for others. Data were shown as mean ± s.d. and compared using a two-sided Student’s t-test; \*\*\*\**P*<0.0001; \*\*\**P*<0.001; n.s., not significant. (**E**) Purified GFP-tagged VPS13C-FL only localizes to giant unilamellar vesicles (GUVs) composed in part of dioleoylglycerol (DOG) lipids, which induce packing defects. In contrast, VPS13C-ΔC (lacking the ATG2_C motif and PH domain) does not localize to GUVs, even in the presence of DOG lipids. (**F**) Overexpressed mCherry-tagged ATG2_C region in the cell lysate only localizes to GUVs with DOG lipids. (**G**) Graphic summary of VPS13C regulation. At the C-terminus, the VAB domain covers the end of the bridge domain, where lipids would be transferred to membrane, blocking lipid transfer. Upon lysosomal membrane damage, amphipathic helices in the ATG2_C motif (which are mobile in the soluble autoinhibited state, as indicated by the “fuzzy” red color) associate with membrane, and the Rab7A-VAB interaction further stabilizes VPS13C-membrane attachment. Conformational changes in the VAB domain expose the lipid bridge for transfer. At the N-terminus, CaM binds constitutively and may modulate bridge architecture and/or ER recruitment in response to calcium. Phosphorylation at both termini (red stars) by kinases in calcium signaling pathways may further regulate VPS13C conformation and its activity at contact sites.

We found that Rab7(Q67L) specifically retains full-length VPS13C, although with relatively weak affinity as the retained VPS13C represented only a small fraction (∼5%) of total overexpressed VPS13. The interaction depends on the presence of VPS13C’s VAB domain as retention of a VPS13C construct lacking the VAB was drastically reduced. Removing other C-terminal modules--including the amphipathic helices in the ATG2_C motif, the WWE domain, or the PH domain--or combinations of them did not abrogate VPS13C pull-down, indicating that these regions are not required for interactions with Rab7. In fact, constructs lacking the ATG2_C motif (constructs ΔC and ΔATG2_C) were retained more efficiently by Rab7(Q67L), suggesting a potential inhibition of the binding to Rab7 by this domain (Fig. 4B left two panels), in agreement with previous studies suggesting an inhibitory role of C-terminal domains on Rab7 binding (Wang et al., 2025). A construct encoding the VAB domain alone was retained by Rab7, demonstrating sufficiency of the VAB domain for Rab7 binding (Fig. 4B, right panel). We were also able to isolate a complex between Rab7 and VPS13C using purified proteins, in the presence of cross-linker as the interaction is weak (Suppl. Fig S6), supporting that the interaction is direct (but see below).

Amphipathic helices are well known to insert into lipid mono-and bilayers (Gimenez-Andres et al., 2018), particularly those with packing defects as might be found in damaged lysosomes (Gahlot et al., 2024). Moreover, previous *in cellulo* studies suggested that amphipathic helices in VPS13’s ATG2_C motif act to sense lysosome damage and bind to their membranes (Wang et al., 2025). To complement these earlier studies, we carried out liposome-based assays with purified proteins. In agreement with the cellular studies, we found that full length VPS13C associated with liposomes (84.8% POPC, 10% POPE, 5% DGS-NTA, 0.2% Rhodamine-PE) in flotation assays (Fig 4 C-D), whereas the association of a VPS13C construct lacking the amphipathic helices of the ATG2_C motif, VPS13CΔATG2C, was strongly reduced (∼70% reduction). Binding was not entirely eliminated, likely because other elements at both the N- and C-termini of VPS13CΔATG2C also interact with membranes. VPS13CΔATG2C’s association with liposomes was rescued by tethering purified Rab7(Q67L) to the liposomes via a hexahistidine tag bound to DGS-NTA lipids in the liposomes (Fig. 4D). Consistent with a role for the VAB domain in interactions with Rab7, the presence of Rab7 did not rescue liposome association of another VPS13C construct that additionally lacked the VAB domain (VPS13CΔVABΔATG2C) (Fig. 4D).Importantly, as these experiments were carried out with purified proteins rather than with cell lysates, they provided strong support that the interaction between Rab7 and VPS13C is direct, involves the VAB domain, and occurs in the context of membrane.

In a separate experiment, we monitored the association of purified fluorescently tagged constructs of VPS13C with giant unilamellar vesicles (GUVs) by confocal microscopy. Full-length VPS13C did not associate with POPC-GUVs (89.5% POPC, 10% DOPS, 0.5% Cy5-PE), but only with POPC+DOG-GUVs (59.5% POPC, 10% DOPS, 30% dioleoylglycerol (DOG), 0.5% Cy5-PE), in which DOG introduces packing defects due to its smaller head group (Vamparys et al., 2013). Further, even binding to GUVs containing DOG was no longer observed when a C-terminally truncated VPS13C lacking the ATG2_C region and PH domain (DC_GFP_) was used (Fig. 4E), supporting that the ATG2_C region is required for membrane binding. We next aimed to demonstrate sufficiency of the ATG2_C region for membrane binding. As the ATG2_C fragment of VPS13C could not be purified due to aggregation, we expressed it as an mCherry-tagged construct in Expi293 cells and incubated lysates from these cells with GUVs, finding that colocalization of the protein with GUVs occurs only with GUVs containing DOG (Fig. 4F). Th experiments showed that VPS13C can sense the difference between membranes with and without packing defects, supporting the importance of lysosome damage in the recruitment of VPS13C. Note that full-length VPS13C behaves differently in the GUV-based assays (where binding requires packing defects induced by DOG) (Fig. 4E) versus in the flotation assay (where binding does not require DOG) (Fig. 4C-D). This may be due to the high curvature of the liposomes used for the flotation assay, as high curvature will decrease the packing of phospholipid head groups, or to the different lipid compositions of the liposomes used for the two assays.

Collectively, our in vitro experiments are consistent with and expand upon the conclusions from experiments in cells, that VPS13C localizes to damaged lysosomes via coincidence detection: the amphipathic helices sense membranes with packing defects, and the VAB domain interacts with Rab7 on lysosomal membranes.

## Discussion

Formation and molecular composition of membrane contacts within cells are dynamically regulated in response to cellular signals. It is increasingly appreciated that BLTPs are key players at these sites via their lipid transfer function, but how their localization, assembly into complexes, and function is controlled is still far from understood. Our work on one such protein, VPS13C, has uncovered new aspects of its regulation, and of VPS13 family protein regulation more broadly (Fig. 4G).

Previous studies in cells had suggested that VPS13C might exist in different conformations, as this would explain why a construct encoding only its VAB domain binds lysosomes constitutively, while binding to these organelles of full-length VPS13C is highly regulated (Wang et al., 2025). More specifically, it was shown that VPS13C is present in the cytosol in an autoinhibited state from which it is released when the C-terminal region of VPS13C interacts with damaged lysosomal membranes. One major outcome of our cryo-EM study was to identify an inactive conformation for VPS13C which likely corresponds to this autoinhibited state. We found that when VPS13C is in solution, interactions with the C-terminal “cap” maintain the arch-shaped VAB domain over the end of the bridge domain. In this position, the VAB blocks access of the bridge domain to the acceptor bilayer, thus preventing lipid transfer. We also identified “cap” helix H_3394-3418_ as partially blocking the end of the lipid transfer channel in this lipid-transfer nonpermissive state. Thus, lipid transfer requires conformational rearrangements at the VPS13 C-terminal end, where “cap” helix H_3394-3418_ moves out of the lipid transfer channel and interactions between the “cap” and VAB are loosened, allowing the VAB to move to the side of the bridge and thus to remove the occlusion of its end. Indeed, in a recent cryo-EM study of VPS13A complexed with a receptor (the scramblase XK) at the acceptor membrane (in Hu et al, in preparation), VPS13A is in such a transfer permissive conformation. It is likely that the existence of lipid transfer inactive and active conformations is a feature of all VPS13 proteins as they mostly share the same domain architecture, and thus likely share regulatory mechanisms.

Our present and past (Wang et al., 2025) results suggest that the coincident insertion of the amphipathic helices of the ATG_C motif into the lipid packing defective membrane of the damaged lysosome and association of the VAB domain with Rab7 could trigger the conformational transition between lipid transfer inactive and active forms for VPS13C. A model was proposed in which in the cytosolic (soluble) form of VPS13C (Wang et al., 2025), access of the VAB domain to Rab7 is prevented by C-terminal modules. But the validity of this model remained to be assessed by biochemical and structural studies. Our work here reveals that conformational changes are required not only for Rab7 to access its binding site on the VAB domain but also to shift VPS13 from a lipid transport nonpermissive to a lipid transport permissive state (as in Fig. 4G). There likely are additional triggers – besides ATG_C binding to the lipid acceptor membrane -- for the conformational changes that are not yet identified. For example, we noted before that in the reconstruction, the end of H_3731-3745_ at the very C-terminus of VPS13C butts against a hydrophobic patch on beta-sandwich repeat #1 of the VAB, and we find intriguing that two residues at the end of the helix (S3742, S3743) are reported to undergo phosphorylation (Phosphosite.org). Plausibly their phosphorylation could contribute to precipitating the VAB’s rearrangement relative to the bridge.

A second major outcome of our study is identification of calmodulin as an interactor for the subset of human VPS13s (VPS13A, VPS13C, VPS13D, but not VPS13B) that localize to contacts involving the ER as the lipid donor membrane. We found that CaM also interacts at an analogous site with fungal Vps13, although the predominant physiological binding partner of yeast Vps13s may be another member of the calmodulin superfamily, Cdc31 (De et al., 2017; Kilmartin, 2003), rather than CaM itself. Thus, the function of calmodulin proteins in VPS13-mediated lipid transport is conserved across species, from yeasts to mammals. The binding site for CaM in VPS13 is close to the N-terminal end of its bridge domain, proximal to the long “FFAT”-motif (residues 877-883 in VPS13C) containing loop that mediates binding of VPS13 to the ER protein VAP (Kumar et al., 2018). As the association of the N-terminus of the bridge domain of VPS13 to the ER is required to allow VPS13 to function in lipid extraction from this organelle, an attractive model is that CaM helps control the connection between the bridge-domain and the ER membrane.

The presence of CaM, which is central to intracellular calcium signaling, in VPS13 complexes converges with other findings implicating VPS13 activity in calcium dependent pathways, either directly or indirectly (Soczewka et al., 2019; Yeshaw et al., 2019). In particular, Soczewka et al. reported that sequestration of CaM could ameliorate a phenotype due to VPS13 deficiency in yeast. Since this rescue occurred in the absence of VPS13, the effect must be indirect, likely due to decreased calcineurin activity, as reported in a follow-up study (Wardaszka et al., 2021). A link between calcium and VPS13 is especially appealing because intracellular membrane contact sites are hubs for calcium signaling. Particularly relevant for VPS13C function at ER-lysosome contacts, calcium release from lysosomes is part of the stress response to their damage (Chen et al., 2024; Lloyd-Evans and Waller-Evans, 2020), but the source of released calcium likely varies for other VPS13s (VPS13A, VPS13D, and fungal Vps13) that localize to contacts with other organelles. It is tempting to speculate that in the VPS13 complex, CaM could confer calcium sensitivity to ER localization or lipid transport activity. Each calcium binding site in CaM’s four EF-hands has a different affinity for calcium and calcium binding affects CaM’s conformation and how it interacts with substrate proteins. Interactions with substrate proteins, in turn, affect CaM’s affinity for calcium (Andrews et al., 2020). While CaM associates tightly and likely constitutively with VPS13s in cells regardless of calcium levels, intracellular calcium levels could affect the CaM-VPS13 interface and consequently (i) control VPS13’s ability to associate directly or indirectly with the ER in order to function in lipid transfer or (ii) impact the width of VPS13’s bridge domain, and thus lipid flux.

Many calcium signals are transmitted via post-translational phosphorylation, often involving calcium/calmodulin-dependent kinases (CaMKs). In this context, it is noteworthy that inspection of phosphosite databases reports several sites expected to be controlled by calcium/CaM-dependent kinases (Phosphosite.org), especially at the N- and C-terminal ends of VPS13C which anchor it to donor and acceptor membranes. These include residues S3742 and S3743 referenced above, residue S373 in a loop near the CaM binding site, and residues of the FFAT motif (Phosphosite.org). Phosphorylation of the FFAT motif may enhance VPS13C’s recruitment by VAP on the ER (as (Kors et al., 2022); and, indeed, VPS13D features a phospho-FFAT motif, whose phosphorylation is required for its ER-localization (Guillen-Samander et al., 2021). How post-translational modifications impact VPS13 activity is a fertile ground for future investigation, albeit beyond the scope of the current work.

In conclusion, as exemplified here for VPS13C, the localization of VPS13s and their lipid transfer activity at contact sites involves regulated interaction with protein partners at the interface with membranes; direct interactions with bilayers; regulation by post-translational modifications; and conformational rearrangements (Fig. 4G). In sum, VPS13 activity depends on a highly complex interplay of multiple factors. Our work represents both a significant advance in our understanding of how VPS13s are activated and, moreover, a framework for further research regarding the function of this fascinating class of proteins.

### Data and Code Availability

Cryo-EM maps for VPS13C were deposited at the EMDB, accession numbers EMD-73345 (map #1), EMD-73344 (map #2), EMD-73343 (map #3), EMD-73373 (composite map). Models for VPS13C and its N- and C-terminal ends and for CtVps13α have been deposited at the PDB: accession numbers 9YRP, 9YQP, 9YQQ, 9YRM. All primary and supporting data generated in this study have been deposited in Zenodo (10.5281/zenodo.17417584). Key laboratory materials are provided in Supplementary Table S4.

## Acknowledgments

The authors thank Dr. Joshua Lees for his advice regarding cryo-EM data processing, Drs. Daniel Lorenzo and Stefano Vanni for estimating the number of lipids that fit into the VPS13C bridge domain, Dr. Benjamin Johnson for discussion, and Han Zhou for technical assistance. This work was funded by the NIH (R35GM131715 to KMR and DA018343 to PDC), Aligning Science Across Parkinson’s grant ASAP-000580 through the Michael J. Fox Foundation for Parkinson’s Research to PDC and KMR, the Parkinson Foundation to PDC, and a postdoctoral fellowship from the Kavli Institute for Neuroscience to HH. For the purpose of open access, the author has applied a CC-BY public copyright license to the Author Accepted Manuscript (AAM) version arising from this submission.

## Contributions

KMR & PDC conceived and supervised the project and wrote the manuscript, with input from DL and XW. DL carried out every aspect of the VPS13C structure determination and analysis; built and analyzed the model for *Ct*Vps13α; conceived of, designed and carried out all biochemical experiments shown except those of Table S3 and those involving GUVs. XW designed and carried out GUV-based experiments, performed the “in cellulo” work in Fig. 2G and Fig. S5C and developed the initial protocol for human VPS13C purification. Together with HH he detected CaM in anti-VPS13 immunoprecipitates (Table S3), leading to the identification as CaM of the non-VPS13 density subsequently identified by DL in the cryo-EM maps of CtVps13 and VPS13C. BH shared coordinates for the VPS13A/XK complex before publication and was involved in analysis. SH contributed to Fig 2F, YL to Fig 4B, and GC to Fig. 4E-F.

### Methods

#### DNA Plasmids

All plasmids used in this paper are listed in Table S1. The codon-optimized full-length human VPS13C (UniProt: Q709C8) with a C-terminal 3xFLAG tag, human Rab7A-Q67L (UniProt: P51149) with an N-terminal Strep tag or an N-terminal MBP tag and a C-terminal 6xHis tag, Chaetomium VPS13 (UniProt: G0S3B8) with an N-terminal 3xFLAG tag were cloned into the pCAG vector. VPS13C mutants, ΔATG2C, ΔC, ΔVAB, ΔC+WWE and VAB domain were cloned into the pCAG vector with a C-terminal 3xFLAG tag. Other VPS13C constructs, including FL_GFP_, ΔC_GFP_, and all calmodulin-binding defective mutants were cloned into the pCMV10 vector with a C-terminal GFP tag, followed by 3xFLAG tag. All cloned plasmids were amplified using maxiprep (Catalog # 740414, Takara Bio) before transfection. For bacterial expression, the *Ct*VPS13 fragment (1-635) was cloned into the first multiple cloning site of the pETDuet vector with an N-terminal 6xHis-Strep tag. For co-expression of calmodulin, to the second multiple cloning site in pETDuet, the full-length human calmodulin sequence (*Hs*CaM, UniProt: P0DP23) was inserted with an N-terminal 6xHis tag.

#### Expression and purification of full-length VPS13C and its mutants

Expi293F cells (RRID:CVCL_D615) were cultured in 8% CO2 with constant orbital shaking at 125 rpm, following the manufacturer’s instructions (MAN0007814, Thermo Scientific). For transfection, 200 μg constructs encoding full-length VPS13C or its mutants were transfected using Expifectamine (Cat. #A14525, Gibco) into 200mL Expi293F cells at a density of 2.4-2.8 million cells/ml, and manufacturer supplied enhancers were added 18 hours post-transfection. Cells were harvested after 48 hours of transfection, flash-frozen and stored at -80°C until use. For protein purification, cells were thawed at room temperature and resuspended in protein purification buffer A (500 mM NaCl, 50 mM HEPES, pH 7.8, 10% glycerol, 1 mM TCEP), supplemented with 1x protease inhibitor cocktail (Cat. #11873580001, Roche). Cells were lysed by three rounds of freeze-and-thaw cycles between liquid nitrogen and a water bath at room temperature. The lysate was homogenized in a Dounce homogenizer and then centrifuged at 27,143 rcf for 30 minutes in a JA-20 rotor. The supernatant was incubated with 100 μL anti-FLAG M2 resin (Cat. #A2220, Millipore-Sigma) for 2 hours at 4°C. The resin was washed twice with 15 mL of buffer A, then incubated overnight with buffer A supplemented with 1 mM freshly-made ATP and 2 mM MgCl_2_ to remove chaperones. After two additional washes, proteins were eluted using Buffer A containing 0.25 mg/mL FLAG peptide (Cat. #A6002, Apex Bio). Elution was performed in five sequential 100 µL incubations, each lasting 20-30 minutes. The pooled eluates were loaded onto the Superose 6 10/300 column (Cat. #29091596, Cytiva), pre-equilibrated with buffer B (200 mM NaCl, 50 mM HEPES, pH 7.2, 6% glycerol, 1 mM TCEP). For structural analysis, full-length VPS13C was subjected to size-exclusion chromatography in buffer C (200 mM NaCl, 50mM HEPES, pH 7.2, 1mM TCEP). Peak fractions were collected and concentrated. The detailed protocol for expression, purification, and negative-stain examination of VPS13C and related constructs were deposited in protocol.io (doi.org/10.17504/protocols.io.rm7vz92d4gx1/v1).

#### Negative-stain electron microscopy of full-length VPS13C and its mutants

Negative stain analysis was performed using 400-mesh carbon-coated copper grids (Cat. #CF400-Cu, Electron Microscopy Sciences). Grids were glow-discharged for 35 s at 25 mA in a Sputter Coater (SCD005, Bal-Tec). 5-7 μL of the protein sample, diluted to 50-100 nM, was applied to the grid and incubated for 30 seconds. The sample was blotted and three to five drops of 5 μL of 2% uranyl acetate solution were applied sequentially, with the last drop incubated for 30 seconds before blotting to complete dryness. Images were acquired using the FEI Tecnai T12 transmission electron microscope operated at 120 kV at a nominal magnification of 52,000x, corresponding to 2.14 Å/pixel at the specimen level. 2D classifications of picked particles were performed with CryoSPARC v4.6.2 (Punjani et al., 2017).

#### Cryo-EM sample preparation and data collection

Cryo-EM grids were cleaned by an oxygen/argon plasma with Gatan Solarus Advanced Plasma System (Model 950). A 3.5 μL aliquot of concentrated VPS13C (∼0.04 mg/ml) was applied to Quantifoil R1.2/1.3 300 mesh gold grids coated with a 2nm continuous carbon film (Cat. # 668-300-AU, Ted Pella), and incubated for 30 seconds. Grids were blotted for 1.5 seconds with a blot force of -12 using the Vitrobot Mark IV (Thermo Scientific) under 100% humidity at 8 °C, then plunge-frozen in liquid ethane cooled by liquid nitrogen. Pre-screened grids were stored in liquid nitrogen prior to loading into a Titan Krios transmission electron microscope (FEI) operated at 300 kV, equipped with a BioQuantum energy filter and a K3 direct electron detector (Gatan). Data acquisition was performed using SerialEM (Mastronarde, 2005) in super-resolution mode (0.534 Å/pixel) with a defocus range of -2.0 to -2.5 µm. Each movie was recorded with an exposure time of 2.896 seconds and dose-fractionated into 50 frames, resulting in a total electron dose of 42.7 e⁻/Å². A total of 4,140 movies were collected at 0° tilt and 4,914 movies at 25° tilt. The protocol of VPS13 structural determination has been deposited in protocol.io (doi.org/10.17504/protocols.io.36wgqpqpovk5/v1). Raw movies were deposited into EMPAIR (EMPIAR-13070).

#### Cryo-EM data processing

Cryo-EM data processing was carried out using CryoSPARC v4.6.2 (Punjani et al., 2017). Beam-induced motion was corrected using patch motion correction with a Fourier crop factor of 0.5, resulting in a physical pixel size of 1.068 Å. Contrast transfer function (CTF) parameters were estimated using patch CTF estimation (Punjani et al., 2017). Following manual micrograph curation, 7,178 movies were selected for further processing. Approximately 2.6 million particles were picked using the blob picker with an elliptical blob (100 and 300 Å axis dimensions), and extracted with a box size of 430 pixels (unbinned). After three rounds of 2D classification, 579,426 particles were used to generate six ab-initio models. All particles were subjected to one round of heterogeneous refinement, and the best-resolved volume was further refined using non-uniform refinement. After one round of 3D classification and non-uniform refinement using 392,473 particles, the full-length map (map #1, EMD-73345) has a global resolution of 4.2 Å but the N-terminus remained largely unresolved.

Particles were then re-picked using templates generated from the full-length volume with a 160 Å picking distance, yielding 2,040,737 particles. These particles were extracted with a box size of 512 pixels and down-sampled to 384 pixels. They were split into subsets and mixed with the original 579,426 particles for two rounds of heterogeneous refinement (classification). The highest-resolution class was selected to generate a C-terminal mask. After removing duplicate particles within 100 Å, 403,001 unique particles were subjected to local refinement with re-centering, resulting in a C-terminal map at 3.8 Å resolution (EMD-73344).

A separate round of template picking with a 96 Å separation yielded 4,203,081 particles. They were extracted with a 512-pixel box and down-sampled to 384 pixels. After two rounds of 2D classification, we selected a subset of 190,805 particle that displays improved alignment at the N-terminus. These particles were used for ab initio reconstruction and non-uniform refinement, producing a low-resolution volume representing the N-terminal domain, which was subsequently used to generate a local refinement mask. Simultaneously, the N-terminal volume served as an input volume for two rounds of heterogeneous refinement and one round of 3D classification using all 4,203,081 particles. From this, 461,163 particles were selected and subjected to local refinement with the gaussian prior turned off, yielding an N-terminal map at 4.1 Å resolution (EMD-73343).

All resolutions were estimated using the gold-standard Fourier shell correlation (FSC) 0.143 criterion with noise substitution (Chen et al., 2013; Rosenthal and Henderson, 2003). Final reconstructions for the full-length, C-terminal, and N-terminal maps were post-processed using spIsoNet (Liu et al., 2025) to correct for artifacts caused by preferred particle orientation.

#### Model building and refinement

Initial atomic models corresponding to the C-terminal and N-terminal regions of VPS13C in complex with calmodulin were generated individually using AlphaFold server (Abramson et al., 2024). Structural fragments from these AlphaFold models, including calmodulin, were manually fitted into the cryo-EM density maps in UCSF ChimeraX (Goddard et al., 2018). The models were rebuilt using the DeepMainMast algorithm (Terashi et al., 2024) and RosettaCM (Song et al., 2013). Rebuilt models were fitted into the EM densities using Namdinator (Kidmose et al., 2019). Both N-terminal and C-terminal models were refined using real_space_refine in Phenix with geometry restraints, followed by manual adjustments in Coot and validation in Phenix (Afonine et al., 2018; Casanal et al., 2020). The resulting models were deposited into PDB (PDB-9YQQ; PDB-9YQP).

To generate the full-length model, the N- and C-terminal maps were aligned and combined with the full-length map using the ‘vol max’ command in ChimeraX (Goddard et al., 2018) to generate a composite map (EMD-73373). The final N- and C-terminal models were docked into the composite map. The model for the missing two central RGB domains were generated using AlphaFold and fitted into the composite map. The complete full-length model was refined using Namdinator (Kidmose et al., 2019), manually adjusted in Coot, and further refined and validated in Phenix (Afonine et al., 2018; Casanal et al., 2020) (PDB-9YRP).

For the structure of CtVPS13α (EMDB-21113, (Li et al., 2020)) in complex with calmodulin, an initial model was generated with AlphaFold3 (Abramson et al., 2024) and fitted into the EM density in ChimeraX (Goddard et al., 2018). This model was refined using Namdinator (Kidmose et al., 2019) and Phenix (Afonine et al., 2018), and manually adjusted in Coot (Casanal et al., 2020). Model validation was performed in Phenix (Afonine et al., 2018) (PDB-9YRM). All validation statistics were shown in Table S2. All structure figures were prepared using ChimeraX (Goddard et al., 2018). The process of CryoEM processing and model building have been deposited in protocol.io (doi.org/10.17504/protocols.io.36wgqpqpovk5/v1).

#### Structural comparison of the lipid transfer domain among BLTPs

To compare the architecture of lipid transfer domains across BLTP family proteins, molecular models comprising the N-terminal cap and the first five RBG domains were aligned in ChimeraX with the lipid channel axis facing inward viewing from the N-terminal end. The specific domain boundaries used for alignment were as follows: VPS13A (2-3174), VPS13C (2-3753), *Ct*VPS13 α (2-1231), ATG2A (173-1414; PDB: 8KBY), and BLTP-1 (65-1308; PDB: 9CAP). To enable a focused view of the lipid bridge domain, adaptor domains and flexible loops were removed from all models. Each structure was clipped from the N-terminus using a thin plane perpendicular to the channel axis, guided by the *Side View* module in ChimeraX (Goddard et al., 2018). The camera view and clipping plane were fixed relative to each other to maintain consistent imaging depth and magnification across all sections in every model. Starting from the N-terminus, the clipping plane was moved in 5 Å increments along the channel axis. Each slice was exported as a separate image, normalized to identical dimensions and pixel size, and compiled into animations (Movie S1). To estimate the width and the cross-sectional area of the channel, slice images were analyzed in ImageJ (Schneider et al., 2012). Real-space dimensions were calibrated in ChimeraX (Goddard et al., 2018) using the *Measure and Color Blobs* module on three representative clips. These measurements were then applied to quantify channel geometry across all slices. Data were plotted in GraphPad Prism 6 (RRID:SCR_002798). The alignment of BLTPs’ models and their comparisons were deposited in Zenodo (doi.org/10.5281/zenodo.17417584).

#### Mass spectrometry analysis

Full-length human VPS13C (UniProt Q709C8) fused to a C-terminal GFP-3×FLAG tag was expressed in Expi293F cells (Gibco) and purified as described. The protein was eluted with Buffer A supplemented with 0.25 mg/mL FLAG peptide, and the eluate was analyzed by mass spectrometry at the W. M. Keck Foundation Biotechnology Resource Laboratory, Yale School of Medicine. The list of top 50 proteins co-purified with VPS13C is shown in Table S3.

#### Western blotting

To examine the calmodulin co-purified with full-length VPS13C, VPS13C mutants, and *Ct*VPS13 in various buffer conditions, eluted proteins were resolved on 4–20% Tris-glycine precast gels. Proteins were transferred to PVDF membranes at 90 V for 90 minutes at 4°C in transfer buffer (25 mM Tris, 192 mM glycine, and 20% methanol). Membranes were blocked for 1 hour at room temperature in 3% BSA prepared in Tris-buffered saline containing 0.1% Tween-20 (TBST), then incubated overnight at 4 °C with the anti-calmodulin antibody (Cat. #HY-P82082, MCE), 1:1000 diluted in 3% BSA and TBST. The next day, membranes were washed three times with TBST, incubated for 1 hour at room temperature with the goat anti-rabbit HRP conjugate (Cat. #AP307P, Sigma Aldrich) at 1:1000 dilution. Membranes were washed another three times with TBST and incubated with ECL substrates (Cat. #32106, Thermo Scientific) prior to imaging by ImageQuant LAS 4000.

To confirm the full-length VPS13C and the VAB domain of VPS13C were co-purified with Rab7A(QL), proteins were resolved on the 3–8% Tris-acetate precast gels and subjected to western blot following the above-mentioned procedures. The anti-FLAG primary antibody (Cat. #F1804, Sigma Aldrich) and anti-mouse HRP conjugate (Cat. #62-6520, Thermo Scientific) were both used at 1:1000 dilutions.

To confirm all calmodulin-binding defective mutants have similar expression levels, same quantity of Expi293F cell lysates expressing each mutant was loaded on 3–8% Tris-acetate precast gels and later blot for FLAG (Cat. #F1804, Sigma Aldrich) and α-Tubulin (Cat. #2125, CST) with corresponding secondary HRP conjugates.

To examine the quality of the ATG2_C domain of VPS13C, lysates from Expi293F cells expressing Flag–mCherry–ATG2_CVPS13C were loaded on 4–20% Tris–glycine precast gels (Bio-Rad) and transferred to nitrocellulose membranes. The membranes were probed with anti-FLAG antibody (Cat. #F1804, Sigma Aldrich), followed by secondary antibodies conjugated to IRDye 800CW (LI-COR; 1:10,000), and imaged using the Odyssey imaging system (LI-COR) according to the manufacturer’s instructions. The protocol has been deposited in protocols.io (doi.org/10.17504/protocols.io.kqdg31w17l25/v1).

#### Expression and purification of *Ct*VPS13(1-635)

Both pETDuet-6xHis-Strep-*Ct*VPS13(1-635) and pETDuet-6xHis-Strep-*Ct*VPS13(1-635)-6xHis-*Hs*CaM constructs were transformed to homemade BL21(De3) pLysS cells (Cat. #200132, Agilent). For both constructs, 1 liter of Luria Broth (LB) culture was grown to an OD600 of 0.8, induced with 0.5mM IPTG and harvested after culturing at 18°C for 16 hours. Cells were resuspended in buffer A, supplemented with 5mM imidazole (Cat. #56750, Sigma). The suspension was passed 5 times through Avestin EmulsiFlex-C5 operating at 7,500 psi followed by a centrifugation step at 36,945 rcf for 30 minutes at 4°C. The supernatant was incubated in batch with Talon metal affinity Resin (Cat. #635502, Takara) for 30 minutes while rotating at 4°C. Affinity resin was washed with 50 column volumes of buffer A containing 20mM imidazole and protein was eluted using buffer A containing 300 mM imidazole. The protocol has been deposited in protocols.io (doi.org/10.17504/protocols.io.ewov11m17vr2/v1).

#### Cell culture, transfection and live-cell imaging

HeLa cells (Cat. #RCB5388; RRID: CVCL_R965) were cultured at 37 °C with 5% CO2 in DMEM medium (Cat. #11965092, Gibco) supplemented with 10% FBS (Cat. #A5256701, Gibco). For live-cell imaging experiments, the cells were seeded onto glass-bottomed dishes (MatTek). Following incubation for 24 h, the cells were transfected using FuGene HD (Cat. #E2311, Promega) according to the manufacturer’s recommendations. Spinning-disk confocal imaging was preformed 16–24 h post transfection. Growth media were changed with live-cell imaging solution (Life Technologies) shortly before imaging. Imaging was performed at 37 °C in 5% CO2 using a Nikon Ti2-E inverted microscope equipped with a Spinning Disk Super Resolution by Optical Pixel Reassignment Microscope (Yokogawa CSU-W1 SoRa, Nikon) and Microlens-enhanced Nipkow Disk with pinholes and a ×60 SR Plan Apo IR oil-immersion objective. For the LLOMe experiments, a 2×stock solution was prepared in the imaging solution and added to the dish during imaging. A detailed description of cell culture, transfection and imaging is available at dx.doi.org/10.17504/protocols.io.eq2lyp55mlx9/v1.

#### Strep pull-down of VPS13C mutants by Strep-Rab7A-Q67L

The proteins were co-expressed and purified from Expi293 cells as mentioned above. For co-transfection, 70 μg of plasmid encoding N-terminally Strep-tagged Rab7A and 30 μg plasmid encoding C-terminally FLAG-tagged VPS13C FL or mutants were mixed in Opti-MEM (Cat. #31985070, Gibco), and transfected into 100 mL of Expi293F cells. Following 48 hours of expression, cells were harvested and resuspended in 10 mL buffer A supplemented with 1 mM GTP (Cat. # G8877, Sigma) and 2 mM MgCl_2_, then lysed by three cycles of freeze-thaw. After lysate clarification, the supernatant was divided, with 30% subjected to FLAG pull down using 50 μL anti-Flag M2 resin. The remaining 70% was used for Strep pull down with 50 μL Strep-Tactin XT 4Flow high-capacity resin (Cat. #2-5030-002, IBA) for 2 hours at 4°C. The resin was washed four times with 15 mL buffer A. Elution was performed three times, each in 50 µL buffer A supplemented with 0.5 mg/mL FLAG peptide for FLAG pull down or 50 mM D-Biotin (Cat. #BG-00, G-Biosciences) for Strep pull down and incubating for 20 minutes. Eluates were pooled and concentrated to a final volume of 50 μL. For protein quantification, Eluates from the Strep pull down were analyzed by SDS-PAGE. Rab7A was quantified on 4–20% Tris-glycine precast gels (Cat. #4561096, Bio-Rad). And the co-purified VPS13Cs were resolved on 3–8% Tris-acetate precast gels (Cat. #EA0375BOX, Invitrogen). The amount of co-purified VPS13C was normalized to the quantity of Rab7A to evaluate relative binding affinity among VPS13C mutants. The VAB domain alone of VPS13C that was co-purified with Rab7A cannot be quantified by Coomassie Blue, yet detectable by anti-FLAG western blot. For quantification and comparison, the band intensity of each VPS13C construct retained by Rab7 was normalized by the corresponding Rab7 band intensity, and then normalized to the full-length (FL) construct. Four biological replicates were performed as independent experiments, except ΔC+WWE, where two replicates were performed. Data were plotted as mean ± s.d. and compared using a two-sided Student’s t-test.

#### Cross-linking of VPS13C and Rab7A(Q67L)

The MBP-TEV-Rab7A-6xHis construct was expressed in Expi293F cells and purified following the same protocol used for VPS13C with a few modifications. For MBP-affinity purification, the clarified lysate was incubated with Amylose resin (Cat. # E8021, NEB) for 2 hours at 4°C, washed, and proteins were eluted with buffer A supplemented with 50 mM maltose (M5895, Sigma). Pooled eluates were run into the Superose 6 size exclusion column in buffer B to remove maltose. For cross-linking, purified full-length VPS13C-3xFLAG and MBP-Rab7A-6xHis were mixed at a 1:10 molar ratio, and incubated overnight at 4°C. Cross-linking was initiated by adding 0.1% glutaldehyde to VPS13C alone, Rab7A alone, or the VPS13C+Rab7A mixture, followed by incubation for 15 minutes at room temperature. The reactions were quenched with 100 mM Tris (pH 8.0) for 10 minutes at room temperature. All samples were subject to MBP pull down as described above. The VPS13C cross-linked to Rab7A was analyzed by SDS-PAGE. The detailed protocol of VPS13C-Rab7A complex formation has been deposited in protocols.io (doi.org/10.17504/protocols.io.eq2ly4q4qlx9/v1).

#### Preparation of SUVs

POPC (Cat. #850457), POPE (Cat. #850757), DGS-NTA (Ni) (Cat. #790404), and Lissamine Rhodamine B-PE (Rhod-PE; Cat. #810150) were purchased from Avanti Polar Lipids. Lipids were mixed at a molar ratio of 84.8% POPC, 10% POPE, 5% DGS-NTA and 0.2% Rhod-PE. The mixture was dried under a gentle nitrogen stream and desiccated under vacuum overnight. The dried lipid film was rehydrated in buffer E (200 mM NaCl, 25 mM HEPES, pH 7.5, 1 mM TCEP) to a final concentration of 10 mM. The suspension was incubated at 37°C for 60 minutes with gentle mixing every 20 minutes, followed by five freeze-thaw cycles. The liposomes were then extruded 31 times through a 200 nm polycarbonate filter using the Avanti Mini-Extruder to generate SUVs.

#### Flotation assay of SUVs

In a 100 µL reaction, 1-5 μg purified full-length VPS13C or VPS13C-ΔATG2C was mixed with 1 mM SUVs. For experiments where Rab7A(QL)-6xHis was used, the protein was produced by TEV cleavage of the MBP-TEV-Rab7A-6xHis construct and further purification through size-exclusion chromatography. Alternatively, the Rab7A(QL)-6xHis was produced from bacteria as mentioned above for the *Ct*VPS13(1-635) construct. Equal mass of Rab7A(QL)-6xHis or 6xHis-ATG7 (negative control, gift from the Melia Lab, Yale University) was added with VPS13C-ΔATG2C to SUVs in the presence of 1 mM GTP and 2 mM Mg^2+^. The mixture was incubated at room temperature for 20 minutes.

For the floatation assay, a 90 µL portion of the mixture was combined with equal volume of 60% OptiPrep (Cat. #D1556, Sigma-Aldrich) to generate the 30% layer, and transferred to a 0.8 mL ultracentrifuge tube (Cat. #344090, Beckman). The sample was layered with 300 μL of 20% OptiPrep, followed by 100 μL of reconstitution buffer. The gradient was centrifuged at 193,911xg for 60 minutes in a SW-55 Ti rotor. After centrifugation, 100 μL from the top fraction, which contains all floated SUVs was recovered. A 10 µL input, along with the 30% of the top fraction was analyzed by SDS-PAGE on 3–8% Tris-acetate precast gels and quantified using ImageJ. For quantification, the normalized quantity of proteins in the top fraction was divided by the normalized quantity of proteins from the input, and shown as a percentage. Four biological replicates were performed as independent experiments for FL and ΔATG2C constructs; three biological replicates were performed for other samples. Data were shown as mean ± s.d. and compared using a two-sided Student’s t-test. The detailed protocol of SUV-related experiments has been deposited in protocols.io (doi.org/10.17504/protocols.io.q26g7n9nqlwz/v1).

#### Preparation of giant unilamellar vesicles (GUVs)

GUVs were generated as previously described (ref) with a few modifications. Specifically, coverslips were sonicated in distilled water and ethanol, then incubated at room temperature in 10% APTES ((3-aminopropyl) trimethoxysilane; Sigma, A3648) in ethanol for 30 min. After incubation, coverslips were washed with ethanol and distilled water, followed by incubation in 2.5% glutaraldehyde (GA; Sigma, G6257) in PBS for 30 min. Coverslips were washed in distilled water, dried, and stored at 4 °C. Then, PAA gels were prepared from acrylamide (AA; 40% w/v, Sigma, A4058) and N,N′-methylenebisacrylamide (BAA; 2% w/v, Sigma, M1533) stock solutions. A 1 mL gel solution contained 250 µL acrylamide, 10 µL bis-acrylamide, and 729 µL H_2_O. Solutions were degassed in a vacuum chamber for 30 min before polymerization. Polymerization was initiated by adding 10 µL ammonium persulfate (APS; Sigma) and 1 µL N,N,N′,N′-tetramethylethylenediamine (TEMED; Bio-Rad, 1610801). 30 µL of the solution was placed on a clean glass plate, and the prepared coverslips were placed on top, ensuring full contact. After 15 min of polymerization, coverslips were carefully removed with a razor blade, washed with water, and dried at 50 °C for 30 min. To generate GUVs, all lipids were purchased from Avanti Polar Lipids. Lipid mixtures were dissolved in chloroform at 1 mg/mL in glass vials. The lipid compositions (molar %) were: POPC-GUVs: 89.5% POPC, 10% DOPS (Cat. #840035, Avanti Polar Lipids), 0.5% Rhod-PE or Cy5-DOPE (Cat. #810335, Avanti Polar Lipids). DOG-GUVs: 59.5% POPC, 10% DOPS, 30% DOG (Cat. #800811, Avanti Polar Lipids), 0.5% Rhod-PE or Cy5-DOPE. Coverslips were placed in 6-well plates, and 20 µL of lipid solution was applied onto each gel. A glass Drigalski spatula was used to evenly spread the solution. The solvent was evaporated in a vacuum chamber for 30–40 min. 1 mL of buffer (500 mM sucrose, 25 mM HEPES, pH 7.4) was carefully added to the wells and incubated overnight. Vesicles were collected by gently pipetting the solution, repeatedly aspirating and dispensing, and transferred to clean 1.5 mL tubes.

#### Confocal fluorescence microscopy of protein–GUV association

GUVs (1 µL) were incubated with either 4 µL purified proteins (300 nM working concentration) or cell lysates in 35 mm glass-bottom MatTek dishes (pre-passivated with BSA) at room temperature for 10 min. Images were acquired using the same Spinning Disk Super Resolution by Optical Pixel Reassignment Microscope described above. All images were analyzed with ImageJ. The detailed protocol of GUV-related experiments has been deposited in protocols.io (doi.org/10.17504/protocols.io.81wgbwko1gpk/v1).

#### Data availability

All primary and supporting data generated in this study have been deposited in Zenodo (doi.org/10.5281/zenodo.17417584). Key laboratory materials and protocols are provided in Supplementary Table S4 and in Zenodo (doi.org/10.5281/zenodo.17459272).

---

**Figure S1.**
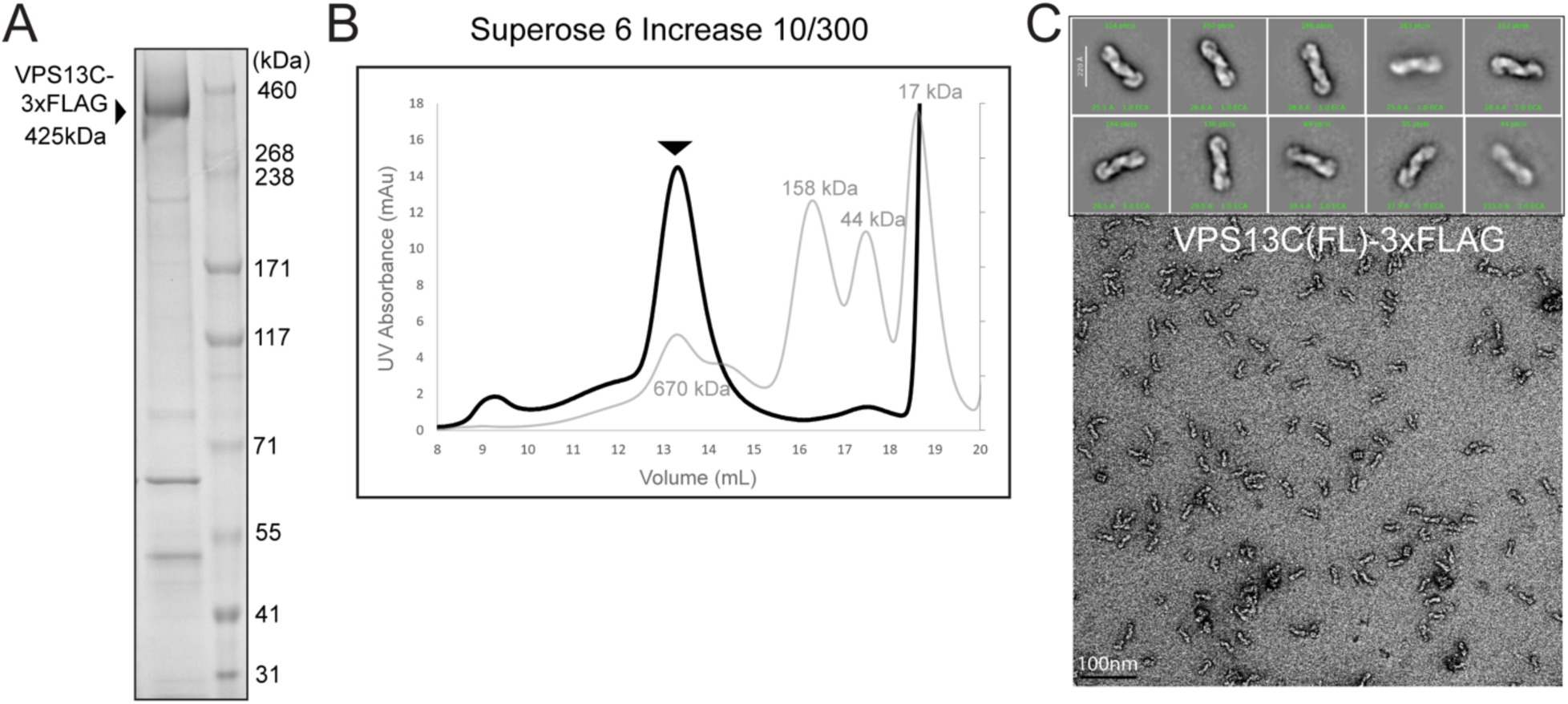
Preparation of full-length VPS13C for Cryo-EM studies. (A) Purified full-length VPS13C-3xFLAG expressed in Expi293F cells analyzed by SDS-PAGE. (B) Size-exclusion chromatography of VPS13C-3xFLAG (425 kDa) on a Superose 6 10/300 column shows a monodisperse peak eluting near the 670 kDa standard. (C) Representative negative-stain EM image and 2D averages reveal a homogeneous sample with a rod-like shape.

**Figure S2.**
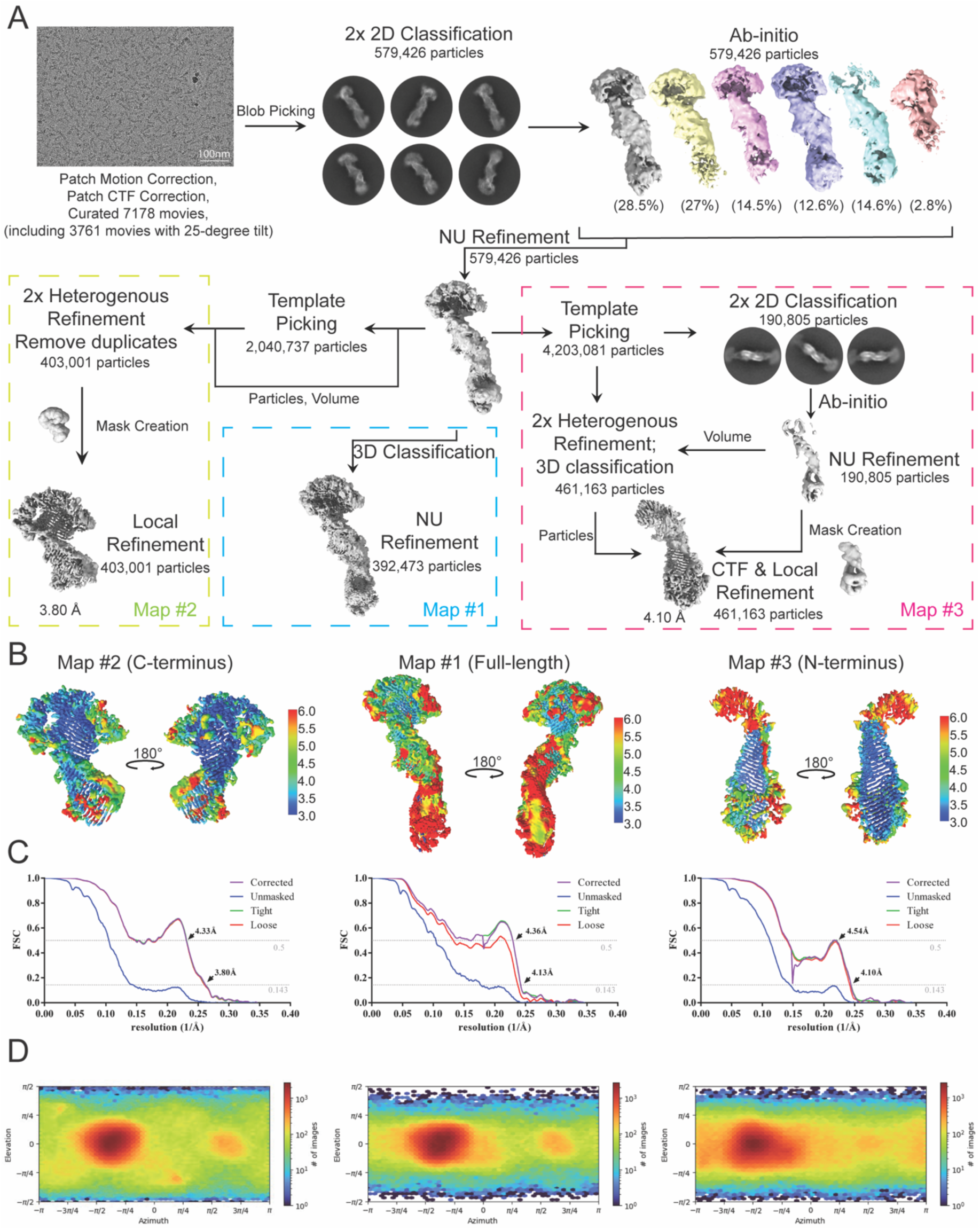
Data processing of full-length VPS13C-3xFLAG. (A) Workflow of data processing. (B) Local resolution estimation of the three final maps, shown from front and back views. Map #1 is the lower-resolution full-length map; Map #2 and Map #3 are locally-refined maps of the C-terminus and the N-terminus, respectively.(C) FSC curves from final refinements for all three maps. (D) Orientation distribution plots for particles contributing to the three maps, showing preferred orientation.

**Figure S3.**
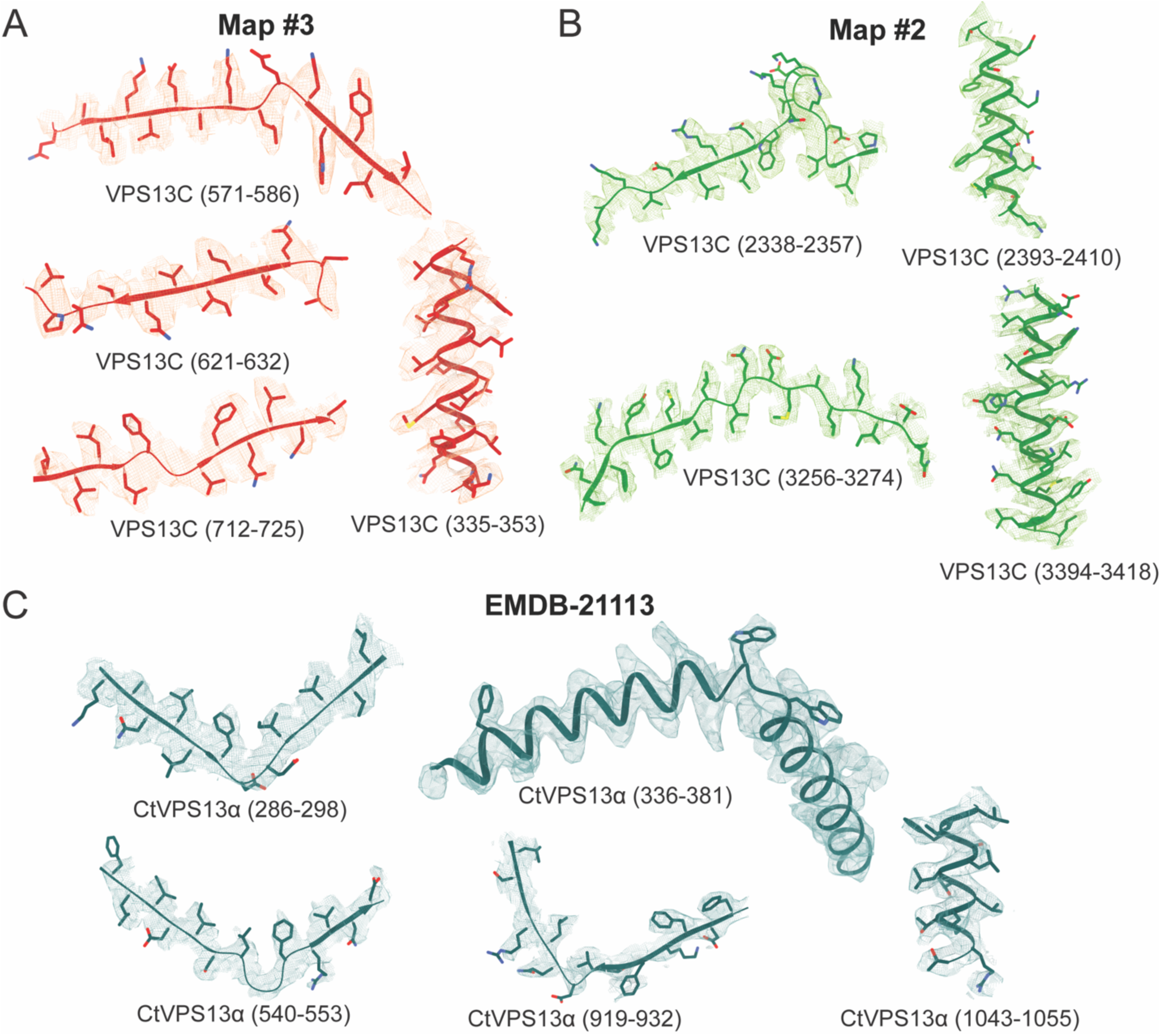
Examples of model fitting into EM maps for VPS13C and *Ct*VPS13α. (A) Representative map and model fits for regions in the VPS13C’s N-terminus. (B) Representative map and model fits for regions in the VPS13C’s C-terminus.(C) Representative map and model fits for regions in *Ct*VPS13α.

**Figure S4.**
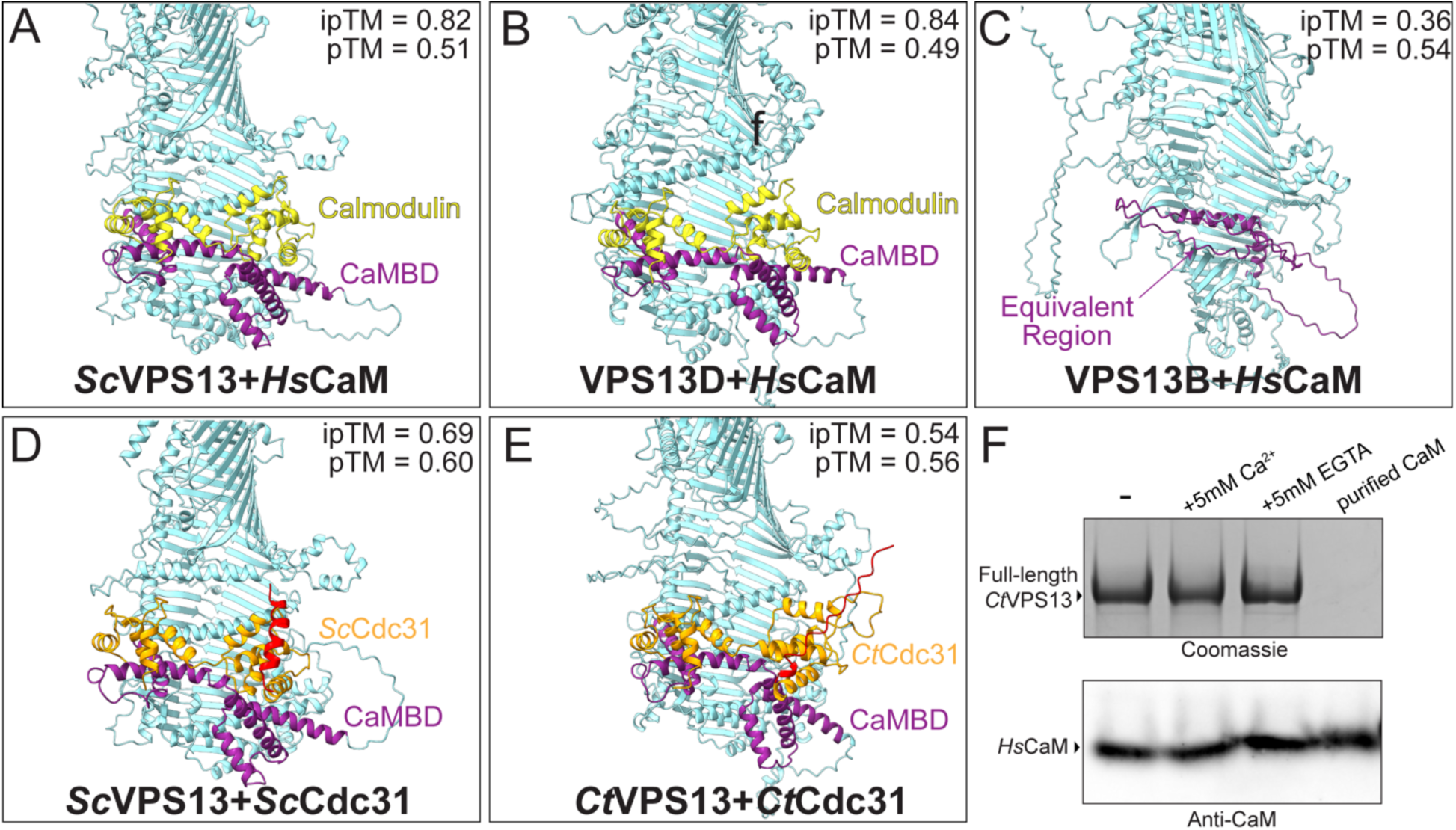
AlphaFold3 predicts interactions between the calmodulin superfamily and VPS13 proteins. (A) Predicted interaction of Saccharomyces cerevisiae (*Sc*) VPS13 (light blue) with CaM (yellow) through a conserved CaM-binding domain (CaMBD, purple). (B) Predicted interaction of human VPS13D with CaM, colored as in (A). (C) Prediction of human VPS13B and CaM. The CaMBD-equivalent region between RBG_1_ and RBG_2_, is highlighted in purple. CaM is not predicted to interact with any part of VPS13B shown here. (D) Predicted interaction of *Sc*VPS13 with *Sc*Cdc31. *Sc*Cdc31 (orange) differs from human CaM by an additional N-terminal helix (red). (E) Predicted interaction of *Ct*VPS13 with *Ct*Cdc31. Similarly, *Ct*Cdc31 (orange) differs from human CaM by an additional N-terminal loop (red). (F) When overexpressed in Expi293F cells, *Ct*VPS13 co-purifies with CaM in FLAG-IP. The binding of calmodulin to *Ct*VPS13 is unaffected by the presence of calcium or EGTA throughout the purification. Because of the difference in sizes for *Ct*VPS13 and CaM, the same samples were run twice on different gels (4-20% Tris-glycine and 3-8% Tris-acetate) to best resolve both proteins.

**Figure S5.**
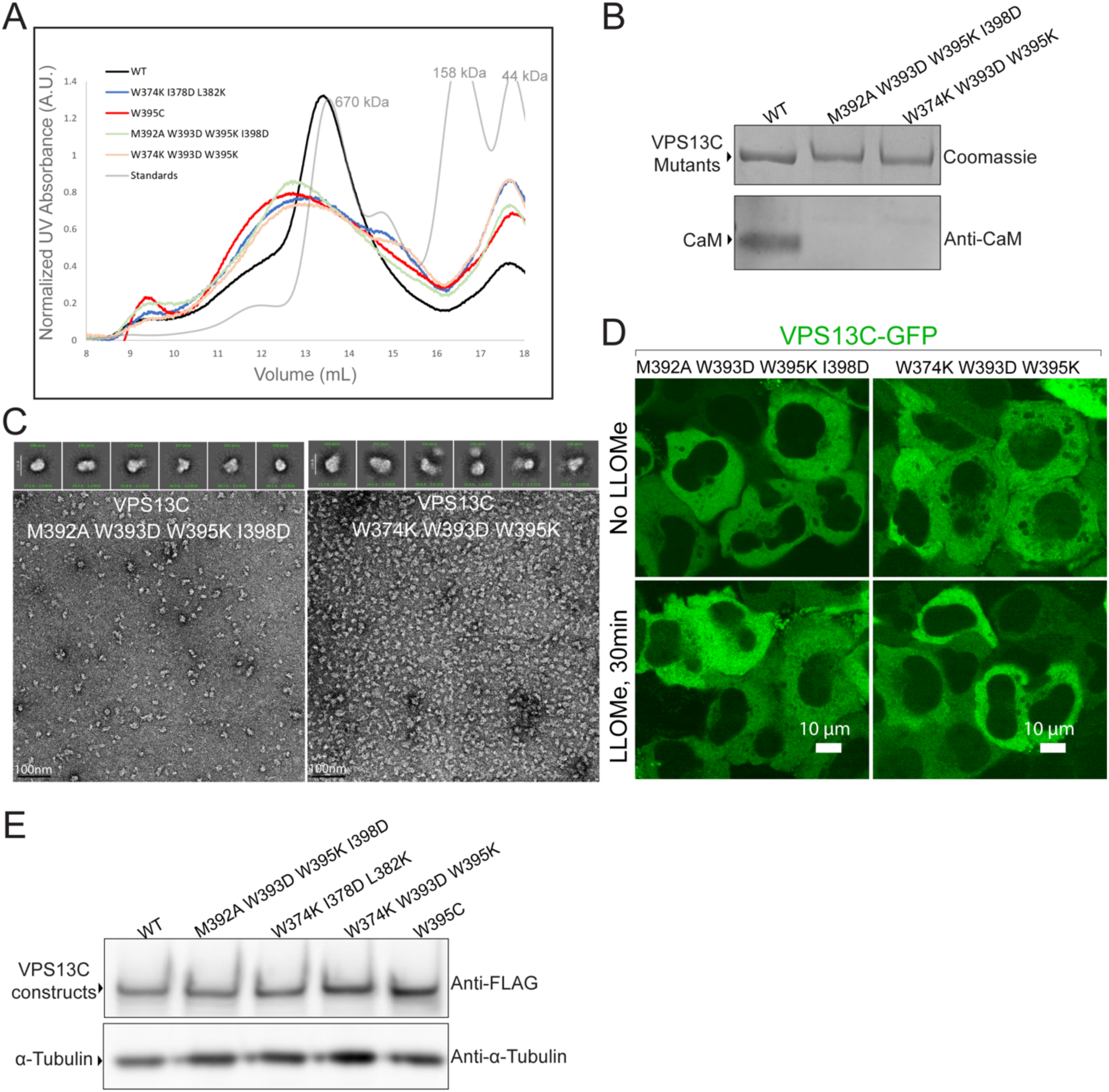
Characterization of calmodulin-binding defective VPS13C mutants. (**A**) Size-exclusion chromatography of VPS13C WT and calmodulin-binding defective mutants. Mutants elute with broader and shifted peaks compared to WT, yet remain soluble and do not aggregate in the void volume. (**B**) Like W395C, additional mutants M392A/W393D/W395K/I398D and W374K/W393D/W395K do not bind CaM, (**C**) lose the rod-like shape of WT protein but remain not non-aggregated, (**D**) and fail to localize to membrane contact sites upon LLOME treatment. (**E**) All CaM-binding defective mutants are expressed at levels comparable to VPS13C WT when overexpressed in Expi293F cells.

**Figure S6.**
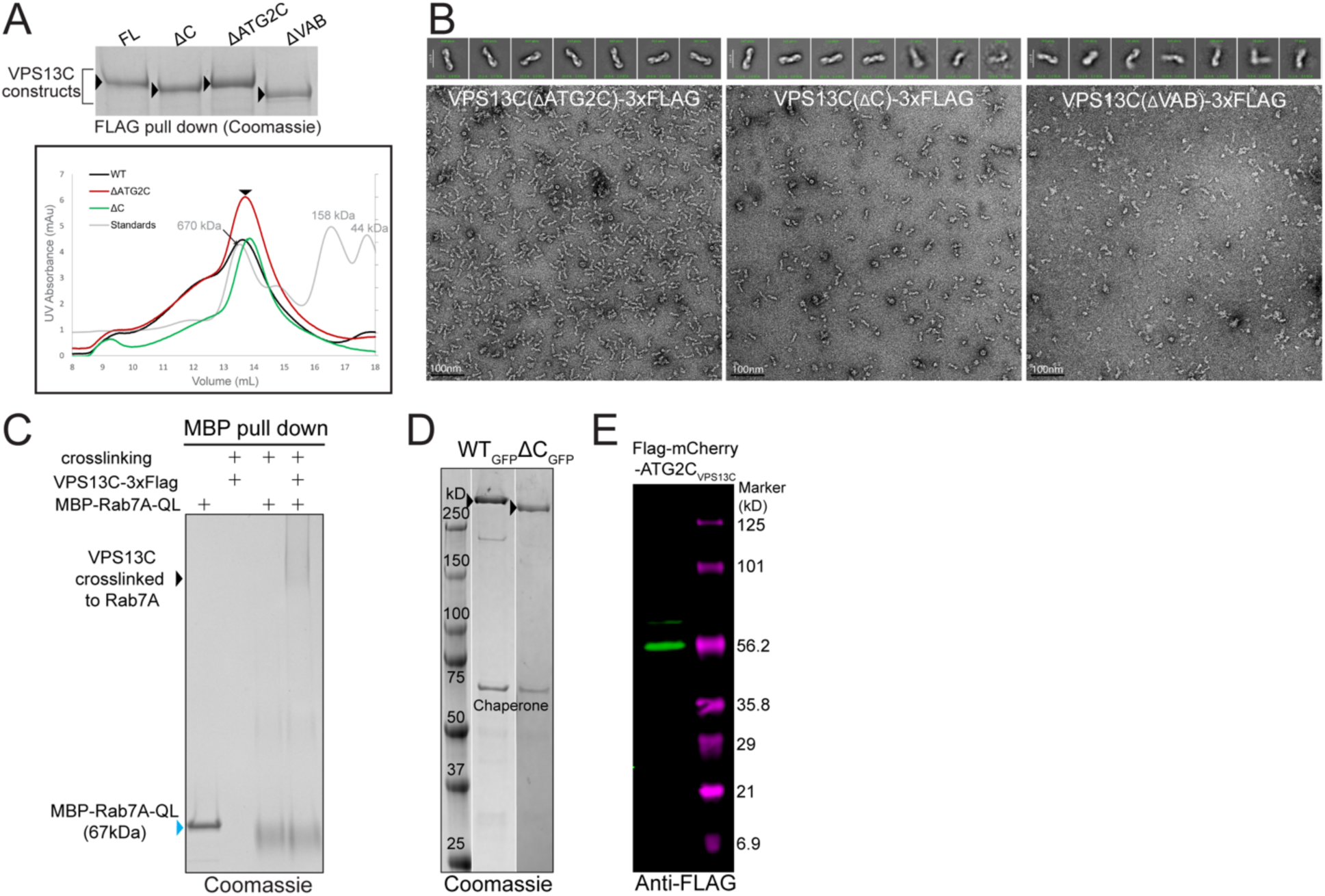
Quality control of VPS13C constructs by purification, EM, and crosslinking. (**A**) VPS13C deletion constructs used for Rab7A pull down experiments and liposome flotation assays are not degraded or aggregated, and migrate at their expected molecular weights on the gel and by size-exclusion chromatography. (**B**) Representative negative-stain EM images and 2D averages show that VPS13C deletion constructs are well-folded and not aggregated. (**C**) Purified full-length FLAG-tagged VPS13C forms a complex with MBP-tagged Rab7A upon glutaldehyde crosslinking. (**D**) GFP-tagged VPS13C WT and VPS13C-ΔC used for GUV experiments are pure and intact, not degraded. (**E)** mCherry-tagged VPS13C-ATG2C used for GUV experiments is intact in cell lysate.

**Supplementary Table S1.**
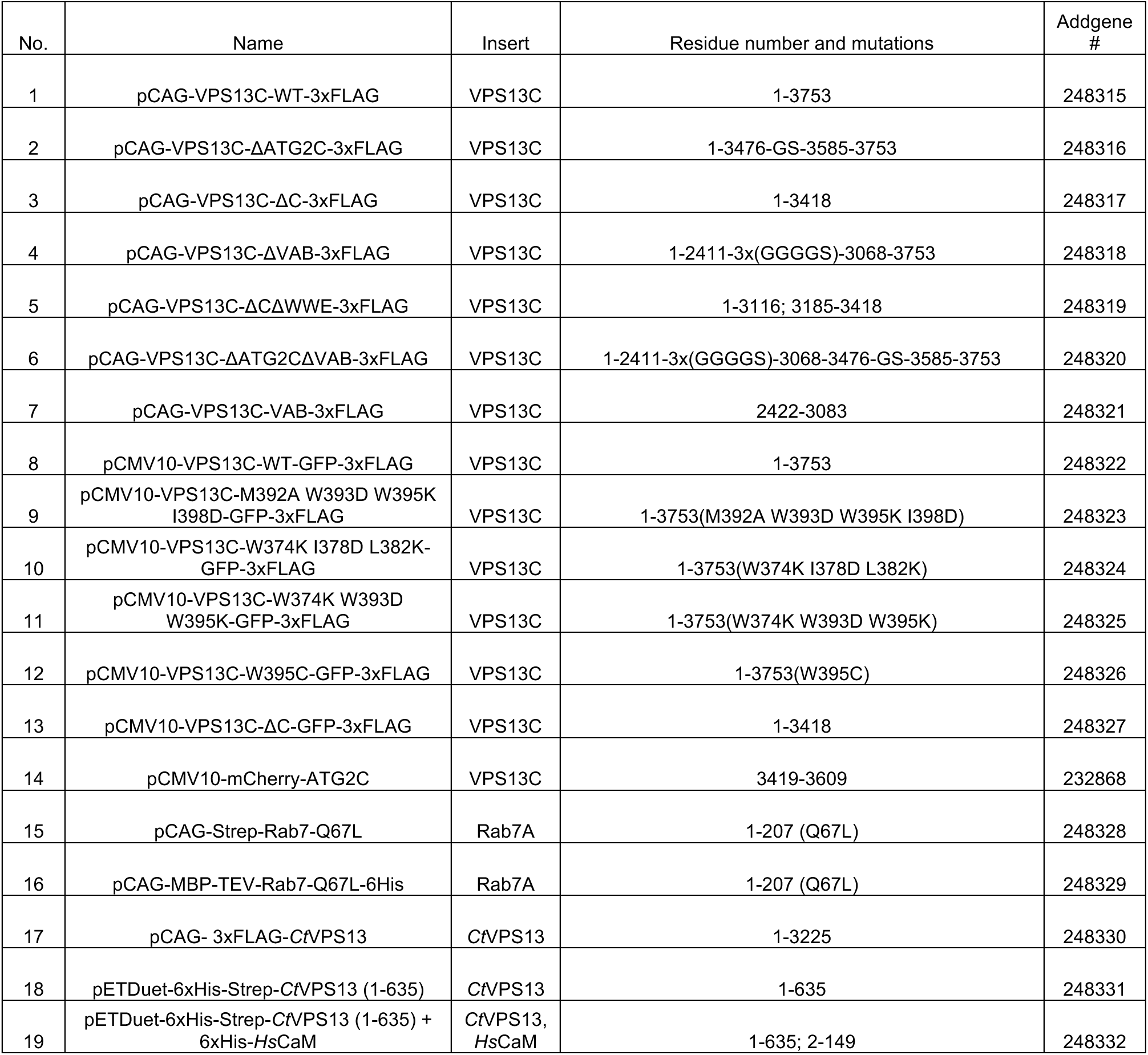
Plasmids used in this study.

**Supplementary Table S2.**
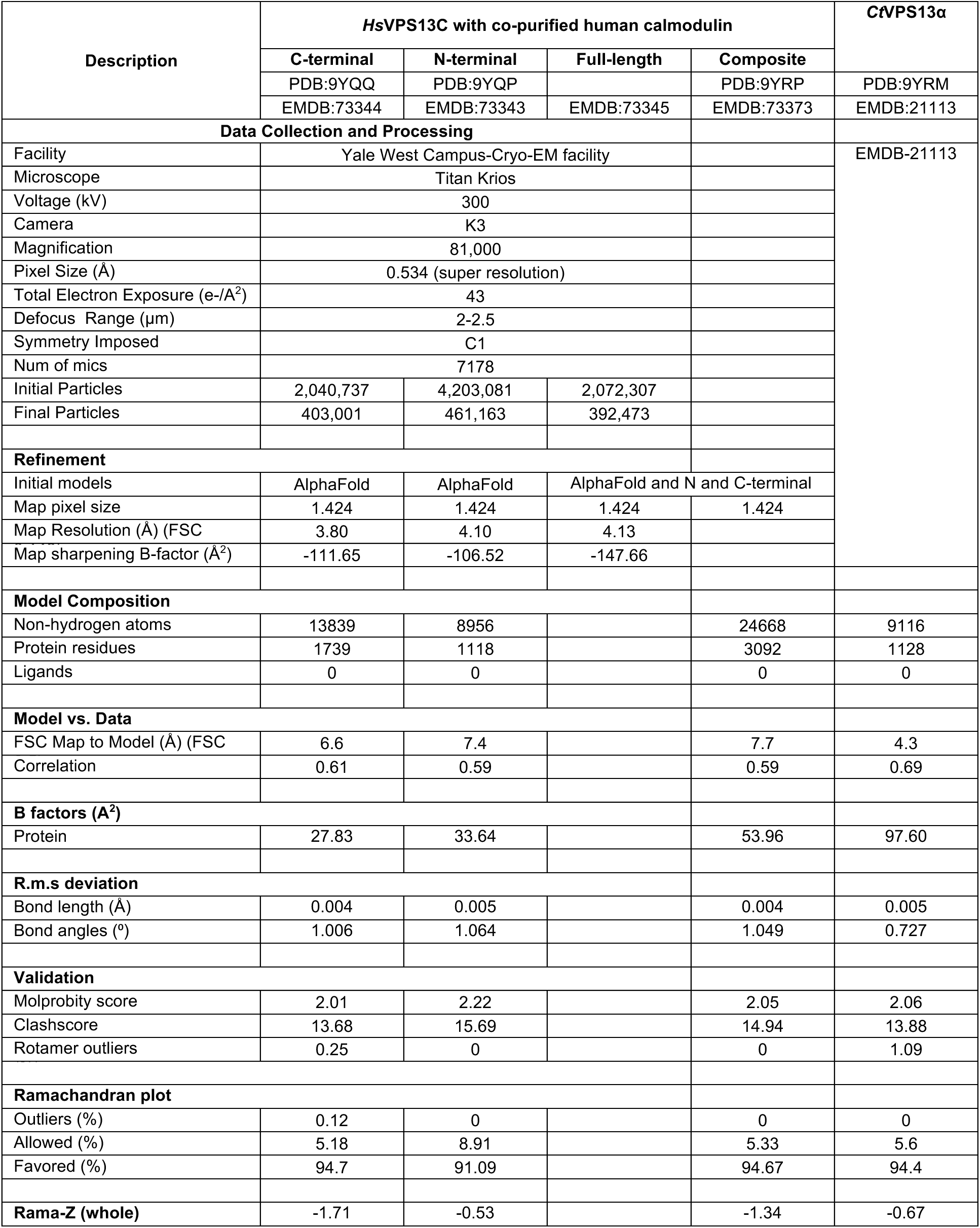
Cryo-EM and model statistics.

| Description | HsVPS13C with co-purified human calmodulin |  |  |  | CtVPS13α |
| --- | --- | --- | --- | --- | --- |
|  | C-terminal | N-terminal | Full-length | Composite |  |
|  | PDB:9YQQ | PDB:9YQP |  | PDB:9YRP | PDB:9YRM |
|  | EMDB:73344 | EMDB:73343 | EMDB:73345 | EMDB:73373 | EMDB:21113 |
| Data Collection and Processing |  |  |  |  |  |
| Facility | Yale West Campus-Cryo-EM facility |  |  |  | EMDB-21113 |
| Microscope | Titan Krios |  |  |  |  |
| Voltage (kV) | 300 |  |  |  |  |
| Camera | K3 |  |  |  |  |
| Magnification | 81,000 |  |  |  |  |
| Pixel Size (Å) | 0.534 (super resolution) |  |  |  |  |
| Total Electron Exposure (e-/Å²) | 43 |  |  |  |  |
| Defocus Range (µm) | 2-2.5 |  |  |  |  |
| Symmetry Imposed | C1 |  |  |  |  |
| Num of mics | 7178 |  |  |  |  |
| Initial Particles | 2,040,737 | 4,203,081 | 2,072,307 |  |  |
| Final Particles | 403,001 | 461,163 | 392,473 |  |  |
| Refinement |  |  |  |  |  |
| Initial models | AlphaFold | AlphaFold | AlphaFold and N and C-terminal |  |  |
| Map pixel size | 1.424 | 1.424 | 1.424 | 1.424 |  |
| Map Resolution (Å) (FSC | 3.80 | 4.10 | 4.13 |  |  |
| Map sharpening B-factor (Å²) | -111.65 | -106.52 | -147.66 |  |  |
| Model Composition |  |  |  |  |  |
| Non-hydrogen atoms | 13839 | 8956 |  | 24668 | 9116 |
| Protein residues | 1739 | 1118 |  | 3092 | 1128 |
| Ligands | 0 | 0 |  | 0 | 0 |
| Model vs. Data |  |  |  |  |  |
| FSC Map to Model (Å) (FSC | 6.6 | 7.4 |  | 7.7 | 4.3 |
| Correlation | 0.61 | 0.59 |  | 0.59 | 0.69 |
| B factors (Å²) |  |  |  |  |  |
| Protein | 27.83 | 33.64 |  | 53.96 | 97.60 |
| R.m.s deviation |  |  |  |  |  |
| Bond length (Å) | 0.004 | 0.005 |  | 0.004 | 0.005 |
| Bond angles (°) | 1.006 | 1.064 |  | 1.049 | 0.727 |
| Validation |  |  |  |  |  |
| Molprobity score | 2.01 | 2.22 |  | 2.05 | 2.06 |
| Clashscore | 13.68 | 15.69 |  | 14.94 | 13.88 |
| Rotamer outliers | 0.25 | 0 |  | 0 | 1.09 |
| Ramachandran plot |  |  |  |  |  |
| Outliers (%) | 0.12 | 0 |  | 0 | 0 |
| Allowed (%) | 5.18 | 8.91 |  | 5.33 | 5.6 |
| Favored (%) | 94.7 | 91.09 |  | 94.67 | 94.4 |
| Rama-Z (whole) | -1.71 | -0.53 |  | -1.34 | -0.67 |

**Supplementary Table S3:** Top hits that co-purified with VPS13C.

| # | Identified Proteins (361) | Accession Number | Alternate ID | Molecular Weight |
| --- | --- | --- | --- | --- |
| 1 | Cluster of VPS13C-linker-GFP-linker-3xFlag | VPS13C-linker-GFP-linker-3xFlag [3] |  | 454 kDa |
| 2 | Heat shock 70 kDa protein 1A OS=Homo sapiens OX=9606 GN=HSPA1A PE=1 SV=1 | P0DMV8 | HSPA1A | 70 kDa |
| 3 | Cluster of Tubulin alpha-1A chain OS=Homo sapiens OX=9606 GN=TUBA1A PE=1 SV=1 (Q71U36) | Q71U36 [3] | TUBA1A | 50 kDa |
| 4 | Cluster of Tubulin beta chain OS=Homo sapiens OX=9606 GN=TUBB PE=1 SV=2 (P07437) | P07437 [6] | TUBB | 50 kDa |
| 5 | Cluster of Calmodulin-1 OS=Homo sapiens OX=9606 GN=CALM1 PE=1 SV=1 (P0DP23) | P0DP23 [2] | CALM1 | 17 kDa |
| 6 | Microtubule-associated protein 1B OS=Homo sapiens OX=9606 GN=MAP1B PE=1 SV=2 | P46821 | MAP1B | 271 kDa |
| 7 | Cluster of Heat shock cognate 71 kDa protein OS=Homo sapiens OX=9606 GN=HSPA8 PE=1 SV=1 (P11142) | P11142 [3] | HSPA8 | 71 kDa |
| 8 | Serine/threonine-protein kinase 38 OS=Homo sapiens OX=9606 GN=STK38 PE=1 SV=1 | Q15208 | STK38 | 54 kDa |
| 9 | Cluster of Ras-related protein Rab-1A OS=Homo sapiens OX=9606 GN=RAB1A PE=1 SV=3 (P62820) | P62820 [10] | RAB1A | 23 kDa |
| 10 | Serine/threonine-protein kinase 38-like OS=Homo sapiens OX=9606 GN=STK38L PE=1 SV=3 | Q9Y2H1 | STK38L | 54 kDa |
| 11 | OTU domain-containing protein 4 OS=Homo sapiens OX=9606 GN=OTUD4 PE=1 SV=4 | Q01804 | OTUD4 | 124 kDa |
| 12 | Perilipin-3 OS=Homo sapiens OX=9606 GN=PLIN3 PE=1 SV=3 | O60664 | PLIN3 | 47 kDa |
| 13 | Cluster of Keratin, type I cytoskeletal 10 OS=Homo sapiens OX=9606 GN=KRT10 PE=1 SV=6 (P13645) | P13645 [3] | KRT10 | 59 kDa |
| 14 | Keratin, type II cytoskeletal 1 OS=Homo sapiens OX=9606 GN=KRT1 PE=1 SV=6 | P04264 | KRT1 | 66 kDa |
| 15 | E3 ubiquitin-protein ligase TRIM21 OS=Homo sapiens OX=9606 GN=TRIM21 PE=1 SV=1 | P19474 | TRIM21 | 54 kDa |
| 16 | Cluster of Filamin-A OS=Homo sapiens OX=9606 GN=FLNA PE=1 SV=4 (P21333) | P21333 [2] | FLNA | 281 kDa |
| 17 | Cluster of Ras-related protein Rab-5C OS=Homo sapiens OX=9606 GN=RAB5C PE=1 SV=2 (P51148) | P51148 [3] | RAB5C | 23 kDa |
| 18 | Heterogeneous nuclear ribonucleoprotein M OS=Homo sapiens OX=9606 GN=HNRNPM PE=1 SV=3 | P52272 | HNRNPM | 78 kDa |
| 19 | Tetratricopeptide repeat protein 1 OS=Homo sapiens OX=9606 GN=TTC1 PE=1 SV=1 | Q99614 | TTC1 | 34 kDa |
| 20 | Keratin, type II cytoskeletal 2 epidermal OS=Homo sapiens OX=9606 GN=KRT2 PE=1 SV=2 | P35908 | KRT2 | 65 kDa |
| 21 | Actin, cytoplasmic 1 OS=Homo sapiens OX=9606 GN=ACTB PE=1 SV=1 | P60709 (+1) | ACTB | 42 kDa |
| 22 | Ras-related protein Rab-7a OS=Homo sapiens OX=9606 GN=RAB7A PE=1 SV=1 | P51149 | RAB7A | 23 kDa |
| 23 | Protein arginine N-methyltransferase 5 OS=Homo sapiens OX=9606 GN=PRMT5 PE=1 SV=4 | O14744 | PRMT5 | 73 kDa |
| 24 | Cluster of Keratin, type II cytoskeletal 6B OS=Homo sapiens OX=9606 GN=KRT6B PE=1 SV=5 (P04259) | P04259 [2] | KRT6B | 60 kDa |
| 25 | ADP-ribosylation factor GTPase-activating protein 1 OS=Homo sapiens OX=9606 GN=ARFGAP1 PE=1 SV=2 | Q8N6T3 | ARFGAP1 | 45 kDa |
| 26 | Heat shock protein HSP 90-alpha OS=Homo sapiens OX=9606 GN=HSP90AA1 PE=1 SV=5 | P07900 | HSP90AA1 | 85 kDa |
| 27 | Cluster of Heat shock protein HSP 90-beta OS=Homo sapiens OX=9606 GN=HSP90AB1 PE=1 SV=4 (P08238) | P08238 [2] | HSP90AB1 | 83 kDa |
| 28 | Melanoma-associated antigen D2 OS=Homo sapiens OX=9606 GN=MAGED2 PE=1 SV=2 | Q9UNF1 | MAGED2 | 65 kDa |
| 29 | Exportin-2 OS=Homo sapiens OX=9606 GN=CSE1L PE=1 SV=3 | P55060 | CSE1L | 110 kDa |
| 30 | Polyubiquitin-B OS=Homo sapiens OX=9606 GN=UBB PE=1 SV=1 | P0CG47 (+3) | UBB | 26 kDa |
| 31 | Cluster of Serpin B3 OS=Homo sapiens OX=9606 GN=SERPINB3 PE=1 SV=2 (P29508) | P29508 [2] | SERPINB3 | 45 kDa |
| 32 | Cluster of CLIP-associating protein 2 OS=Homo sapiens OX=9606 GN=CLASP2 PE=1 SV=3 (O75122) | O75122 [2] | CLASP2 | 141 kDa |
| 33 | Protein S100-A9 OS=Homo sapiens OX=9606 GN=S100A9 PE=1 SV=1 | P06702 | S100A9 | 13 kDa |
| 34 | Protein S100-A7 OS=Homo sapiens OX=9606 GN=S100A7 PE=1 SV=4 | P31151 | S100A7 | 11 kDa |
| 35 | Poly(rC)-binding protein 1 OS=Homo sapiens OX=9606 GN=PCBP1 PE=1 SV=2 | Q15365 | PCBP1 | 37 kDa |
| 36 | Ras-related protein Rab-11B OS=Homo sapiens OX=9606 GN=RAB11B PE=1 SV=4 | Q15907 | RAB11B | 24 kDa |
| 37 | Tumor protein D54 OS=Homo sapiens OX=9606 GN=TPD52L2 PE=1 SV=2 | O43399 | TPD52L2 | 22 kDa |
| 38 | BAG family molecular chaperone regulator 2 OS=Homo sapiens OX=9606 GN=BAG2 PE=1 SV=1 | O95816 | BAG2 | 24 kDa |
| 39 | Keratin, type I cytoskeletal 9 OS=Homo sapiens OX=9606 GN=KRT9 PE=1 SV=3 | P35527 | KRT9 | 62 kDa |
| 40 | Kinesin-like protein KIF11 OS=Homo sapiens OX=9606 GN=KIF11 PE=1 SV=2 | P52732 | KIF11 | 119 kDa |
| 41 | Small ribosomal subunit protein eS4, X isoform OS=Homo sapiens OX=9606 GN=RPS4X PE=1 SV=2 | P62701 | RPS4X | 30 kDa |
| 42 | Spartin OS=Homo sapiens OX=9606 GN=SPART PE=1 SV=1 | Q8N0X7 | SPART | 73 kDa |
| 43 | c-Myc-binding protein OS=Homo sapiens OX=9606 GN=MYCBP PE=1 SV=3 | Q99417 | MYCBP | 12 kDa |
| 44 | RuvB-like 2 OS=Homo sapiens OX=9606 GN=RUVBL2 PE=1 SV=3 | Q9Y230 | RUVBL2 | 51 kDa |
| 45 | Exportin-1 OS=Homo sapiens OX=9606 GN=XPO1 PE=1 SV=1 | O14980 | XPO1 | 123 kDa |
| 46 | Protein S100-A8 OS=Homo sapiens OX=9606 GN=S100A8 PE=1 SV=1 | P05109 | S100A8 | 11 kDa |
| 47 | Cluster of 14-3-3 protein gamma OS=Homo sapiens OX=9606 GN=YWHAG PE=1 SV=2 (P61981) | P61981 [4] | YWHAG | 28 kDa |
| 48 | Large ribosomal subunit protein eL24 OS=Homo sapiens OX=9606 GN=RPL24 PE=1 SV=1 | P83731 | RPL24 | 18 kDa |
| 49 | Nucleolar protein 56 OS=Homo sapiens OX=9606 GN=NOP56 PE=1 SV=4 | O00567 | NOP56 | 66 kDa |
| 50 | Large ribosomal subunit protein eL13 OS=Homo sapiens OX=9606 GN=RPL13 PE=1 SV=4 | P26373 | RPL13 | 24 kDa |

**Supplementary Table S4:**
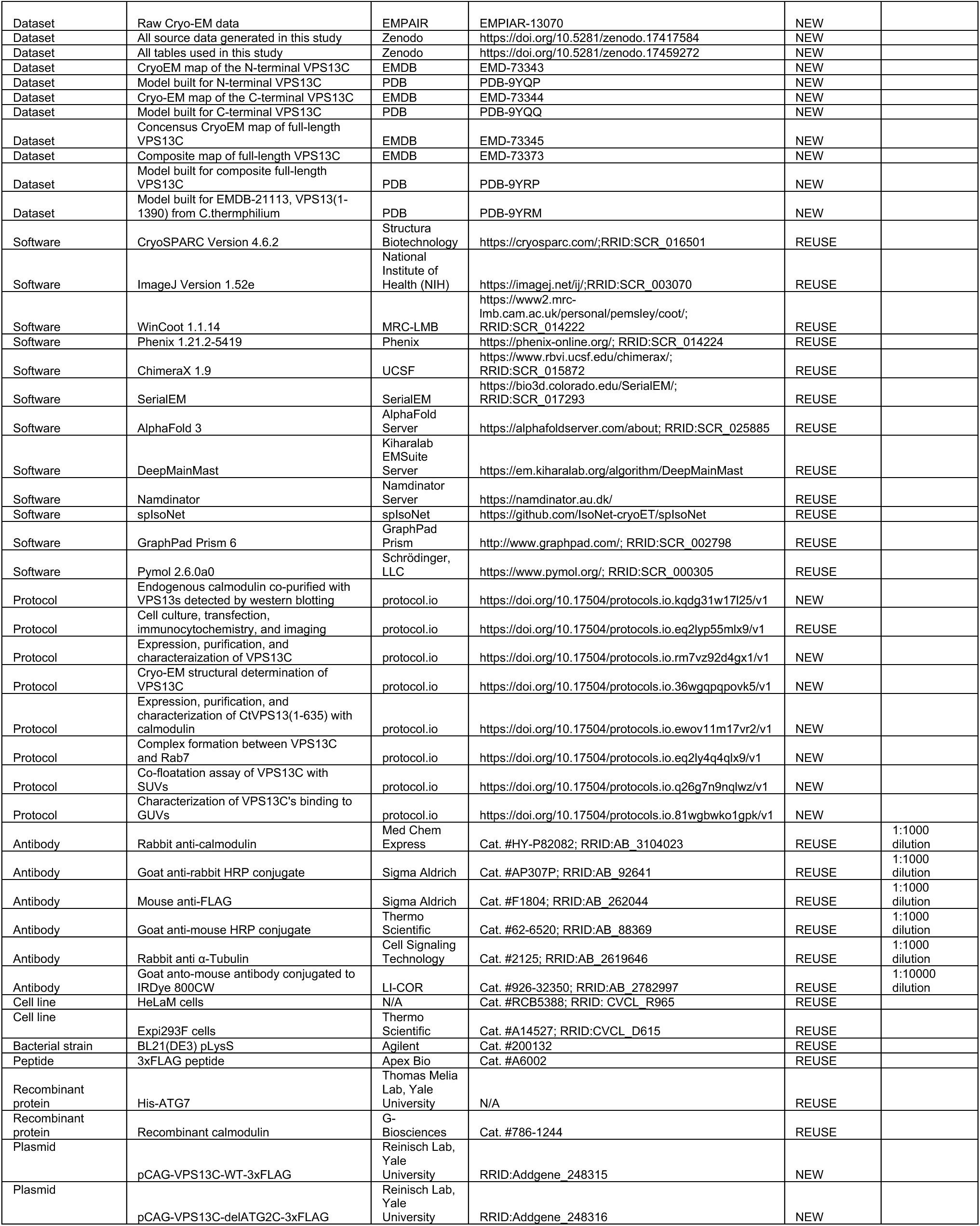
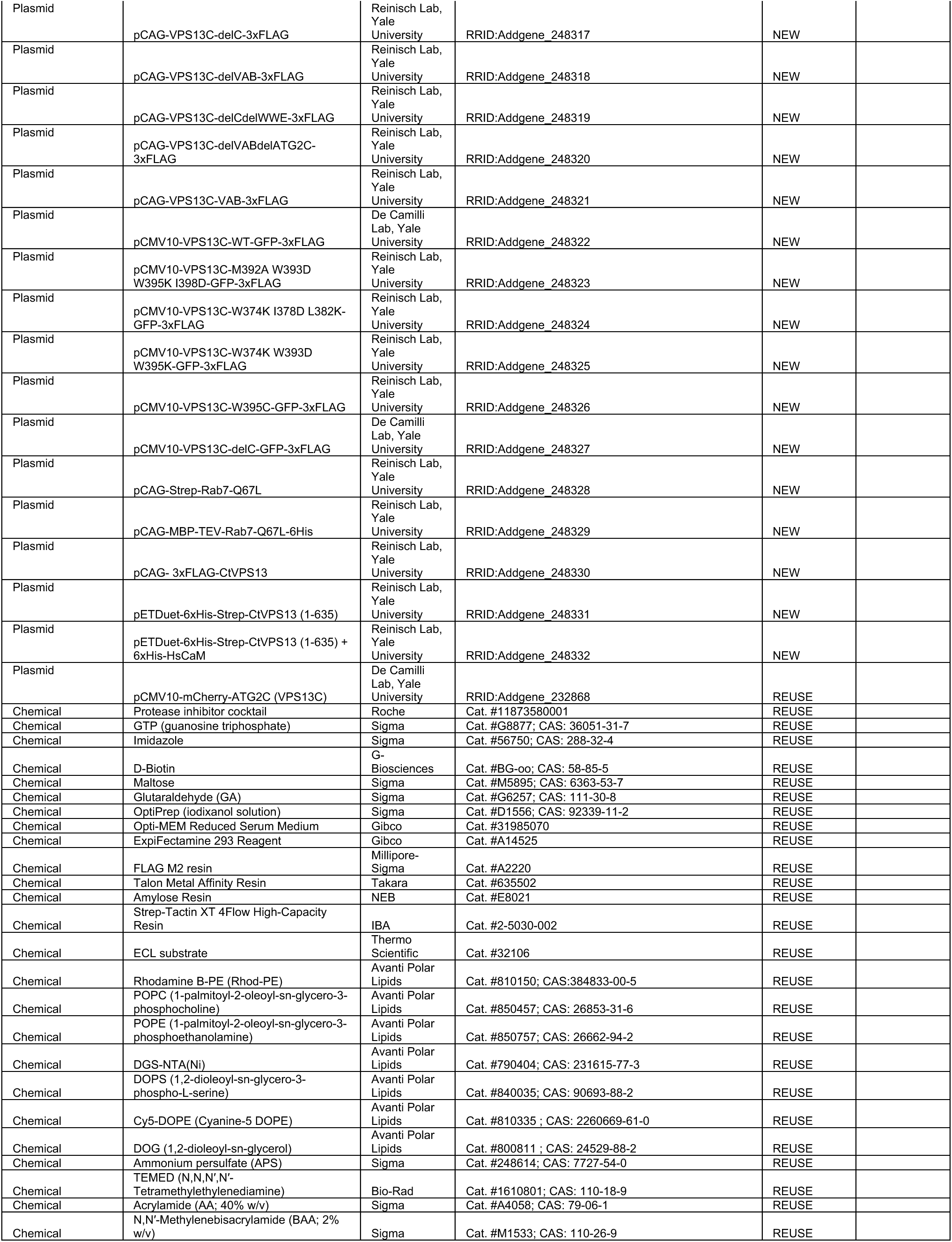
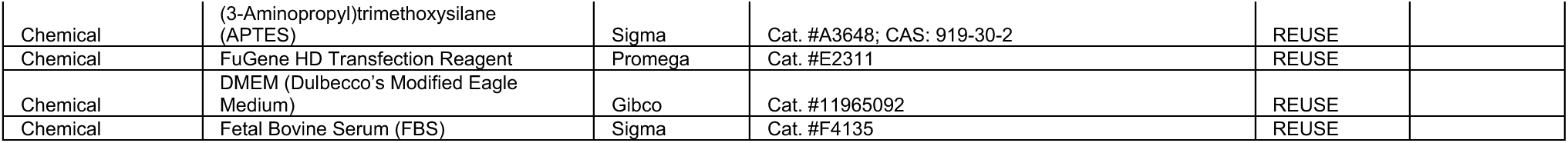
Key lab materials used and generated in this study.

## References

1. Abramson, J., Adler, J., Dunger, J., Evans, R., Green, T., Pritzel, A., Ronneberger, O., Willmore, L., Ballard, A.J., Bambrick, J., et al. (2024). Accurate structure prediction of biomolecular interactions with AlphaFold 3. Nature 630, 493–500.

2. Afonine, P.V., Poon, B.K., Read, R.J., Sobolev, O.V., Terwilliger, T.C., Urzhumtsev, A., and Adams, P.D. (2018). Real-space refinement in PHENIX for cryo-EM and crystallography. Acta Crystallogr D Struct Biol 74, 531–544.

3. Andrews, C., Xu, Y., Kirberger, M., and Yang, J.J. (2020). Structural Aspects and Prediction of Calmodulin-Binding Proteins. Int J Mol Sci 22.

4. Casanal, A., Lohkamp, B., and Emsley, P. (2020). Current developments in Coot for macromolecular model building of Electron Cryo-microscopy and Crystallographic Data. Protein Sci 29, 1069–1078.

5. Chen, S., McMullan, G., Faruqi, A.R., Murshudov, G.N., Short, J.M., Scheres, S.H., and Henderson, R. (2013). High-resolution noise substitution to measure overfitting and validate resolution in 3D structure determination by single particle electron cryomicroscopy. Ultramicroscopy 135, 24–35.

6. Chen, W., Motsinger, M.M., Li, J., Bohannon, K.P., and Hanson, P.I. (2024). Ca(2+)-sensor ALG-2 engages ESCRTs to enhance lysosomal membrane resilience to osmotic stress. Proc Natl Acad Sci U S A 121, e2318412121.

7. De, M., Oleskie, A.N., Ayyash, M., Dutta, S., Mancour, L., Abazeed, M.E., Brace, E.J., Skiniotis, G., and Fuller, R.S. (2017). The Vps13p-Cdc31p complex is directly required for TGN late endosome transport and TGN homotypic fusion. J Cell Biol 216, 425–439.

8. Dziurdzik, S.K., and Conibear, E. (2021). The Vps13 Family of Lipid Transporters and Its Role at Membrane Contact Sites. Int J Mol Sci 22.

9. Gahlot, P., Kravic, B., Rota, G., van den Boom, J., Levantovsky, S., Schulze, N., Maspero, E., Polo, S., Behrends, C., and Meyer, H. (2024). Lysosomal damage sensing and lysophagy initiation by SPG20-ITCH. Mol Cell 84, 1556–1569 e1510.

10. Gillingham, A.K., Bertram, J., Begum, F., and Munro, S. (2019). In vivo identification of GTPase interactors by mitochondrial relocalization and proximity biotinylation. Elife 8.

11. Gimenez-Andres, M., Copic, A., and Antonny, B. (2018). The Many Faces of Amphipathic Helices. Biomolecules 8.

12. Goddard, T.D., Huang, C.C., Meng, E.C., Pettersen, E.F., Couch, G.S., Morris, J.H., and Ferrin, T.E. (2018). UCSF ChimeraX: Meeting modern challenges in visualization and analysis. Protein Sci 27, 14–25.

13. Guillen-Samander, A., Leonzino, M., Hanna, M.G., Tang, N., Shen, H., and De Camilli, P. (2021). VPS13D bridges the ER to mitochondria and peroxisomes via Miro. J Cell Biol 220.

14. Hanna, M., Guillen-Samander, A., and De Camilli, P. (2023). RBG Motif Bridge-Like Lipid Transport Proteins: Structure, Functions, and Open Questions. Annu Rev Cell Dev Biol 39, 409–434.

15. Hu, B., Alvarez, D., Rocha-Roa, C., Guyard, V., Li, D., Wang, X., De Camilli, P., Vanni, S., and Reinisch, K.M. (2025). Molecular insights into bulk lipid transport from structural studies of the bridge-like protein VPS13A complexed with the scramblase XK.

16. Jumper, J., Evans, R., Pritzel, A., Green, T., Figurnov, M., Ronneberger, O., Tunyasuvunakool, K., Bates, R., Zidek, A., Potapenko, A., et al. (2021). Highly accurate protein structure prediction with AlphaFold. Nature 596, 583–589.

17. Kang, Y., Lehmann, K.S., Long, H., Jeherson, A., Purice, M., Freeman, M., and Clark, S. (2025). Structural basis of lipid transfer by a bridge-like lipid-transfer protein. Nature 642, 242–249.

18. Kidmose, R.T., Juhl, J., Nissen, P., Boesen, T., Karlsen, J.L., and Pedersen, B.P. (2019). Namdinator - automatic molecular dynamics flexible fitting of structural models into cryo-EM and crystallography experimental maps. IUCrJ 6, 526–531.

19. Kilmartin, J.V. (2003). Sfi1p has conserved centrin-binding sites and an essential function in budding yeast spindle pole body duplication. J Cell Biol 162, 1211–1221.

20. Kors, S., Costello, J.L., and Schrader, M. (2022). VAP Proteins - From Organelle Tethers to Pathogenic Host Interactors and Their Role in Neuronal Disease. Front Cell Dev Biol 10, 895856.

21. Kumar, N., Leonzino, M., Hancock-Cerutti, W., Horenkamp, F.A., Li, P., Lees, J.A., Wheeler, H., Reinisch, K.M., and De Camilli, P. (2018). VPS13A and VPS13C are lipid transport proteins diherentially localized at ER contact sites. J Cell Biol 217, 3625–3639.

22. Lesage, S., Drouet, V., Majounie, E., Deramecourt, V., Jacoupy, M., Nicolas, A., Cormier-Dequaire, F., Hassoun, S.M., Pujol, C., Ciura, S., et al. (2016). Loss of VPS13C Function in Autosomal-Recessive Parkinsonism Causes Mitochondrial Dysfunction and Increases PINK1/Parkin-Dependent Mitophagy. Am J Hum Genet 98, 500–513.

23. Levine, T.P. (2022). Sequence Analysis and Structural Predictions of Lipid Transfer Bridges in the Repeating Beta Groove (RBG) Superfamily Reveal Past and Present Domain Variations Ahecting Form, Function and Interactions of VPS13, ATG2, SHIP164, Hobbit and Tweek. Contact (Thousand Oaks) 5, 251525642211343.

24. Li, P., Lees, J.A., Lusk, C.P., and Reinisch, K.M. (2020). Cryo-EM reconstruction of a VPS13 fragment reveals a long groove to channel lipids between membranes. J Cell Biol 219.

25. Liu, Y.T., Fan, H., Hu, J.J., and Zhou, Z.H. (2025). Overcoming the preferred-orientation problem in cryo-EM with self-supervised deep learning. Nat Methods 22, 113–123.

26. Lloyd-Evans, E., and Waller-Evans, H. (2020). Lysosomal Ca(2+) Homeostasis and Signaling in Health and Disease. Cold Spring Harb Perspect Biol 12.

27. Mastronarde, D.N. (2005). Automated electron microscope tomography using robust prediction of specimen movements. J Struct Biol 152, 36–51.

28. Punjani, A., Rubinstein, J.L., Fleet, D.J., and Brubaker, M.A. (2017). cryoSPARC: algorithms for rapid unsupervised cryo-EM structure determination. Nat Methods 14, 290–296.

29. Rampoldi, L., Dobson-Stone, C., Rubio, J.P., Danek, A., Chalmers, R.M., Wood, N.W., Verellen, C., Ferrer, X., Malandrini, A., Fabrizi, G.M., et al. (2001). A conserved sorting-associated protein is mutant in chorea-acanthocytosis. Nat Genet 28, 119–120.

30. Reinisch, K.M., De Camilli, P., and Melia, T.J. (2025). Lipid Dynamics at Membrane Contact Sites. Annu Rev Biochem 94, 479–502.

31. Rosenthal, P.B., and Henderson, R. (2003). Optimal determination of particle orientation, absolute hand, and contrast loss in single-particle electron cryomicroscopy. J Mol Biol 333, 721–745.

32. Schneider, C.A., Rasband, W.S., and Eliceiri, K.W. (2012). NIH Image to ImageJ: 25 years of image analysis. Nat Methods 9, 671–675.

33. Shen, X., Valencia, C.A., Gao, W., Cotten, S.W., Dong, B., Huang, B.C., and Liu, R. (2008). Ca(2+)/Calmodulin-binding proteins from the C. elegans proteome. Cell Calcium 43, 444–456.

34. Soczewka, P., Kolakowski, D., Smaczynska-de Rooij, I., Rzepnikowska, W., Ayscough, K.R., Kaminska, J., and Zoladek, T. (2019). Yeast-model-based study identified myosin- and calcium-dependent calmodulin signalling as a potential target for drug intervention in chorea-acanthocytosis. Dis Model Mech 12.

35. Song, Y., DiMaio, F., Wang, R.Y., Kim, D., Miles, C., Brunette, T., Thompson, J., and Baker, D. (2013). High-resolution comparative modeling with RosettaCM. Structure 21, 1735–1742.

36. Terashi, G., Wang, X., Prasad, D., Nakamura, T., and Kihara, D. (2024). DeepMainmast: integrated protocol of protein structure modeling for cryo-EM with deep learning and structure prediction. Nat Methods 21, 122–131.

37. Ueno, S., Maruki, Y., Nakamura, M., Tomemori, Y., Kamae, K., Tanabe, H., Yamashita, Y., Matsuda, S., Kaneko, S., and Sano, A. (2001). The gene encoding a newly discovered protein, chorein, is mutated in chorea-acanthocytosis. Nat Genet 28, 121–122.

38. Vamparys, L., Gautier, R., Vanni, S., Bennett, W.F., Tieleman, D.P., Antonny, B., Etchebest, C., and Fuchs, P.F. (2013). Conical lipids in flat bilayers induce packing defects similar to that induced by positive curvature. Biophys J 104, 585–593.

39. Wang, X., Xu, P., Bentley-DeSousa, A., Hancock-Cerutti, W., Cai, S., Johnson, B.T., Tonelli, F., Shao, L., Talaia, G., Alessi, D.R., et al. (2025). The bridge-like lipid transport protein VPS13C/PARK23 mediates ER-lysosome contacts following lysosome damage. Nat Cell Biol 27, 776–789.

40. Wang, Y., Dahmane, S., Ti, R., Mai, X., Zhu, L., Carlson, L.A., and Stjepanovic, G. (2024). Structural basis for lipid transfer by the ATG2A-ATG9A complex. Nat Struct Mol Biol.

41. Wardaszka, P., Soczewka, P., Sienko, M., Zoladek, T., and Kaminska, J. (2021). Partial Inhibition of Calcineurin Activity by Rcn2 as a Potential Remedy for Vps13 Deficiency. Int J Mol Sci 22.

42. Yan, C., Wu, F., Jernigan, R.L., Dobbs, D., and Honavar, V. (2008). Characterization of protein-protein interfaces. Protein J 27, 59–70.

43. Yeshaw, W.M., van der Zwaag, M., Pinto, F., Lahaye, L.L., Faber, A.I., Gomez-Sanchez, R., Dolga, A.M., Poland, C., Monaco, A.P., van, I.S.C., et al. (2019). Human VPS13A is associated with multiple organelles and influences mitochondrial morphology and lipid droplet motility. Elife 8.

